# Independent expansion, selection and hypervariability of the *TBC1D3* gene family in humans

**DOI:** 10.1101/2024.03.12.584650

**Authors:** Xavi Guitart, David Porubsky, DongAhn Yoo, Max L. Dougherty, Philip C. Dishuck, Katherine M. Munson, Alexandra P. Lewis, Kendra Hoekzema, Jordan Knuth, Stephen Chang, Tomi Pastinen, Evan E. Eichler

## Abstract

*TBC1D3* is a primate-specific gene family that has expanded in the human lineage and has been implicated in neuronal progenitor proliferation and expansion of the frontal cortex. The gene family and its expression have been challenging to investigate because it is embedded in high-identity and highly variable segmental duplications. We sequenced and assembled the gene family using long-read sequencing data from 34 humans and 11 nonhuman primate species. Our analysis shows that this particular gene family has independently duplicated in at least five primate lineages, and the duplicated loci are enriched at sites of large-scale chromosomal rearrangements on chromosome 17. We find that most humans vary along two *TBC1D3* clusters where human haplotypes are highly variable in copy number, differing by as many as 20 copies, and structure (structural heterozygosity 90%). We also show evidence of positive selection, as well as a significant change in the predicted human TBC1D3 protein sequence. Lastly, we find that, despite multiple duplications, human *TBC1D3* expression is limited to a subset of copies and, most notably, from a single paralog group: *TBC1D3-CDKL*. These observations may help explain why a gene potentially important in cortical development can be so variable in the human population.

## INTRODUCTION

Gene duplication followed by adaptation is one of the primary forces by which new genes emerge within species (Ohno, 1970). Many of these evolutionary events occur in segmental duplications (SDs), genomic units that are at least one kilobase pair in length and whose duplications are 90% or more identical to one another (Bailey and Eichler 2006). Many human-specific genes reside in SDs, which often continue to vary structurally in our lineage (Bitar et al. 2019). Since the initial publication of the human and chimpanzee genomes, investigations of human-specific SD genes have found that they most often are implicated in xenobiotic recognition, metabolism, immunity, and neuronal development, playing an important role in the evolution of our species (Perry et al. 2007; Kalebic et al. 2008; Dennis et al. 2012).

*TBC1D3* is a primate-specific SD gene family (Paulding et al. 2003). This gene family is dispersed across the two arms of chromosome 17, though most copies in humans map to two expansion blocks at locus chromosome 17q12 (Fig. 1a). Expression data in humans from the Genome-Tissue Expression (GTEx) project reveals *TBC1D3* is modestly expressed globally, with increased expression in testis and brain tissue (GTEx Consortium 2020). *TBC1D3* expression and function were initially observed in prostate tumor samples and originally classified as an oncogene (Hodzic et al. 2006). However, in 2016, Ju et al. showed that transgenic overexpression of *TBC1D3* in the developing mouse brain results in a proliferation of outer radial glial cells and a subsequent expansion and folding of the cortex (Ju et al. 2016). Functionally, two hypotheses have been proposed for the role of *TBC1D3* in cell proliferation. The first, brought forward by Wainszelbaum et al., suggests that *TBC1D3* induces cell proliferation by acting on the insulin-like growth factor-1 (IGF) and epidermal growth factor (EGF) pathways (Wainszelbaum et al. 2012; 2008). In contrast, Hou et al. conducted experiments in neuron progenitor and organoid body models and observed that *TBC1D3* antagonizes *G9a* histone methyltransferase, increasing the proliferation of cortical neuron progenitors (Hou et al. 2021).

**Figure 1:**
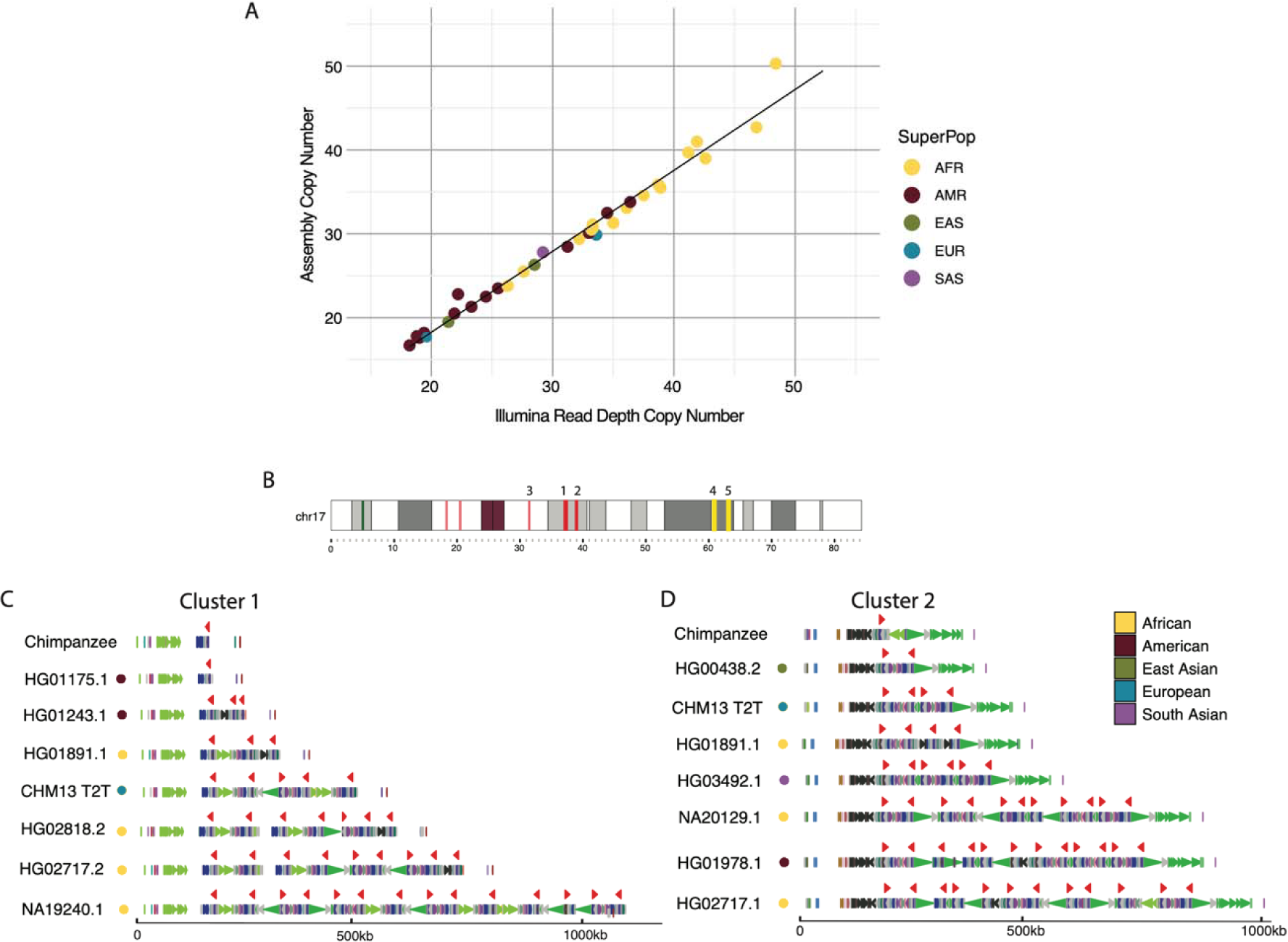
Assembly and human variation of *TBC1D3.* **(A)** Assembly copy number estimate vs. orthogonal Illumina sequence copy number estimate. Each point represents a sample diploid assembly, colored by superpopulation. **(B)** Reference ideogram of *TBC1D3* regions. Expanded views of clusters 1 and 2 are illustrated in **(C** and **D). (C** and **D)** Structure for chimpanzee and seven validated human haplotypes over *TBC1D3* cluster 1 **(C)** and cluster 2 **(D)**. *TBC1D3* copies are colored as red arrows. Colored arrows below *TBC1D3* illustrate segmental duplication content annotated with DupMasker (Jiang et al. 2008).

These findings suggest that the evolution of *TBC1D3* may have contributed to the human cranial expansion over the last two million years (Stringer 2016). Investigations in the sequence evolution and variation amongst humans and nonhuman primates (NHPs) would help test this hypothesis (Hudson, Kreitman, and Aguadé 1987; McDonald and Kreitman 1991; Tajima 1989). However, the duplicated and highly identical sequences of *TBC1D3* copies make assembly impossible with standard short-read sequencing platforms. Instead, researchers have investigated copy number variation in SD genes using short-read sequencing data to understand patterns of variation (Redon et al. 2006). Such read-based studies have suggested extensive copy number differences among human populations (Sudmant et al. 2010). However, these experiments lack the single-base-pair resolution necessary to distinguish different paralogous copies, structural differences among haplotypes, and which copies are likely functional or expressed. Moreover, it is unclear how a gene so variable in copy number could play such a critical role in the expansion of the frontal cortex in humans.

In this study, we address these questions by leveraging long-read sequencing data generated from humans and apes to better understand the variation and evolution of *TBC1D3* (Liao et al. 2023; Mao et al. 2024; Makova et al. 2023). Our analysis shows that this particular gene family has independently duplicated in at least five primate lineages, and that *TBC1D3* duplications are enriched at sites of large-scale chromosomal rearrangements. We find that most human variation maps to two distinct clusters and that humans are highly variable in copy number at these loci, differing by as many as 20 copies and ∼1 Mbp in length depending on haplotypes. Despite this variation, we find that a high level of expression is limited to a subset of *TBC1D3* copies, helping to explain why a gene potentially important in cortical development can be so copy number variable. Furthermore, we find evidence of positive selection and a significant change in the predicted TBC1D3 protein structure in humans, which may drastically alter the function and regulation of TBC1D3.

## RESULTS

### Human *TBC1D3* copy number variation

To understand *TBC1D3* organization and variation in humans, we first focused on two *TBC1D3* gene family clusters that contain the majority of *TBC1D3* paralogs, named cluster 1 and cluster 2 (Fig. 1b). We characterized 44 human genomes recently sequenced as part of the Human Pangenome Reference Consortium at this locus (Liao et al. 2023). We first assessed the integrity of each assembly by searching for sequence collapses in read depth of both Pacific Biosciences (PacBio) high-fidelity (HiFi) and Oxford Nanopore Technologies (ONT) sequencing data (Vollger et al. 2019; Dishuck et al. 2022; Methods). We found that 46 of the haplotypes passed quality control (QC), while 42 haplotypes failed. We attempted to re-assemble the samples that failed QC using a novel assembly algorithm that leverages both HiFi and ONT data (Verkko) (Rautiainen et al. 2023). This procedure recovered an additional 20 haplotypes where both cluster 1 and cluster 2 were fully sequenced and assembled without error (Supplementary Fig. S1). We also confirmed accurate assembly by comparing predicted copy number against Illumina read depth-based copy number estimates, an orthogonal sequencing platform. (Fig. 1a; Methods). In total, we validated 66 haplotypes where both *TBC1D3* clusters were fully resolved and, including three genome references, developed a total dataset of 69 human haplotypes (Supplementary Fig. S10).

Next, we estimated the copy number and organization of *TBC1D3* in clusters 1 and 2 for each human haplotype (Fig. 1b-d). In cluster 1, we found that *TBC1D3* varies from 1 to 14 copies, while in cluster 2 it varies from 2 to 14 copies. The differences in copy account for as much as 1.5 Mbp of differential size between human haplotypes. We find that *TBC1D3* copy number is significantly higher among Africans (X=34.4) when compared to non-African populations (X=25.4) (p-value = 1.7E-5). For cluster 1, we find that 65% (45/69) of the haplotypes are structurally distinct. Interestingly for cluster 2, we observe similar diversity, where 68% (47/69) are structurally distinct (Supplementary Fig. S1). Based on completely assembled diploid samples, we estimate the structural heterozygosity for cluster 1 is 94%, while cluster 2 is 88%, making these two loci among some of the most structurally variable gene families in the human genome (Sudmant et al. 2010).

### Nonhuman primate (NHP) *TBC1D3* organization

To better understand the evolution of the clusters, we investigated the organization of *TBC1D3* in 10 different NHP lineages. This included single representatives of five great ape species (bonobo, chimpanzee, gorilla, Bornean, and Sumatran orangutan), two Old World monkeys (macaque, gelada), two New World monkeys (marmoset, owl monkey), and one prosimian (mouse lemur). Eight of these genomes were previously published (Mao et al. 2024) or are part of efforts to generate telomere-to-telomere (T2T) assemblies of ape genomes (Makova et al. 2023; Data Access). We generated HiFi sequence data from both the gelada and mouse lemur genomes in this study and assembled their genomes using Hifiasm (Methods).

With the exception of the mouse lemur, all NHP genomes carry multiple copies of *TBC1D3*. We find that *TBC1D3* is also highly copy number variable among NHPs, from two copies in marmoset to 31 copies within a single haplotype in both gelada and gibbon. We searched specifically for clustered expansions and found that most primates—human, gorilla, orangutan, macaque, and gelada—similarly contain two expanded clusters of *TBC1D3* (Fig. 2a). Among apes, these two clusters are orthologous to human clusters 1 and 2, separated by 1.35 Mbp of intervening sequence. Among the Old World monkeys, gelada and macaque, structural rearrangements have repositioned the two clusters such that the intervening sequence is larger and nonsyntenic. Importantly, bonobo and chimpanzee only possess 1-2 copies of *TBC1D3* at cluster 2, whereas no copies were identified at cluster 1. Thus, all humans have an increase in copy number when compared to the *Pan* lineage but are not exceptional when compared to most other NHP lineages. New World monkeys, owl monkey, and marmoset do not have *TBC1D3* organized into clusters. Instead, marmoset has two copies while owl monkey has eight copies distributed throughout its chromosome, suggesting independent and recent expansions. Overall, we find that *TBC1D3* copy number varies from 0 to 14 copies in cluster 1 and from 1 to 17 copies in cluster 2 (Fig. 2b). A detailed analysis of the composition of the SDs within each primate lineage shows that the units of duplication in different species frequently differed in structure, suggesting independent duplications or gene conversion events in each lineage (Supplementary Fig. S4; Methods).

**Figure 2:**
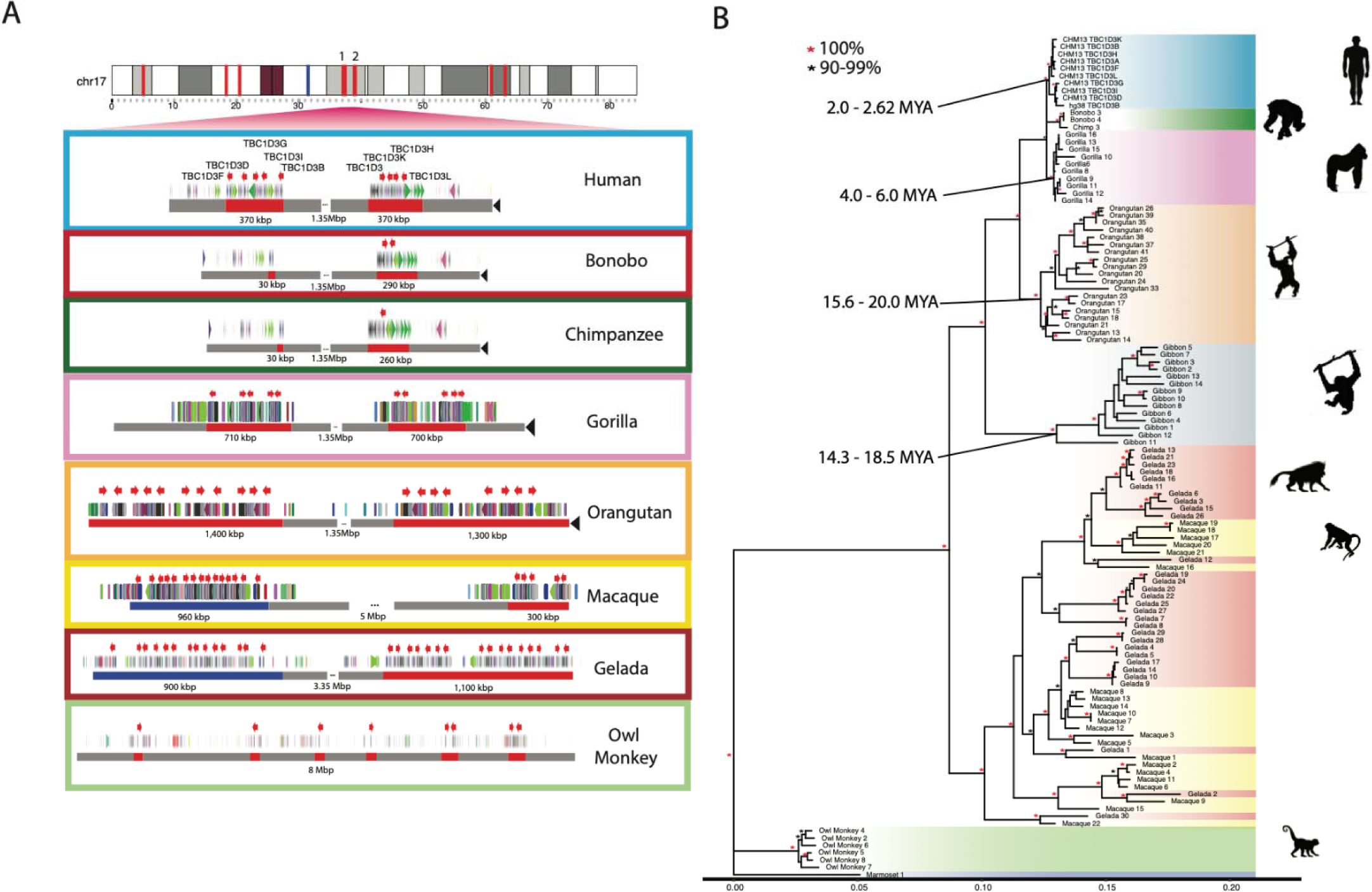
Comparative genome structure and phylogeny of *TBC1D3* gene family among primates. **(A)** *TBC1D3* clusters 1 and 2 structure. Orthologous *TBC1D3* clusters 1 and 2 are illustrated as two clustered regions (red blocks), with flanking unique sequence in gray for the primate lineages. Old World monkey *TBC1D3* expansion 1, which is nonsyntenic, is highlighted with blue blocks. *TBC1D3* paralogs (red arrows) are embedded within other segmental duplication blocks, with DupMasker annotations illustrated with colored arrows. The diverse organizational differences of each expansion, including expansion size, duplicon content, and copy number, suggest independent expansion. **(B)** *TBC1D3* neutral phylogeny generated by maximum likelihood. 2,300 bp of intronic sequence were aligned between all primate *TBC1D3* paralogs observed in (A), with the marmoset sequence used as an outgroup. The phylogeny supports the hypothesis of independent expansion with the exception of the Old World monkeys (gelada and macaque) where several copies duplicated before and after speciation of these two lineages (11 mya) (Liedigk et al. 2014).

In order to estimate when the clustered *TBC1D3* copies expanded in each lineage, we constructed a maximum likelihood phylogenetic tree based on a multiple sequence alignment (MSA) generated from intronic sequence of each predicted *TBC1D3* gene copy from the various primate genomes (Fig. 2b; Methods). We observe complete lineage-specific stratification of the *TBC1D3* gene family members into distinct clades for human, *Pan*, gorilla, orangutan, gibbon, and owl monkey lineages. These findings strongly support recurrent duplication or gene conversion of all gene family copies in each lineage. In contrast, the gelada and rhesus macaque show both shared and lineage-specific groups, suggesting *TBC1D3* expanded before and after speciation. Using 25 and 6.5 million years ago (mya) as times of human–macaque and human–chimpanzee divergence, we estimated the timing of each lineage-specific expansion (Fig. 2b; Stevens et al. 2013; Dunsworth et al. 2010). In most lineages, the primate duplications occurred relatively recently. Most notably, we observe that humans experienced the most recent expansion within the apes, occurring between 2.0 and 2.6 mya.

### *TBC1D3* and large-scale chromosomal rearrangements

During our comparative analysis of NHP genomes, we noticed that chromosomal synteny frequently was disrupted at sites corresponding to interspersed *TBC1D3* loci. To assess this more systematically, we selected five primate lineages where T2T assemblies had recently been generated as part of the Primate T2T Consortium, aligned orthologous chromosome 17s to one another, and illustrated these alignments, as well as alpha satellite and *TBC1D3* loci (Fig. 3a; Methods). We found that *TBC1D3* consistently flanks some of the largest chromosomal rearrangements. For example, human *TBC1D3P2* demarcates one end of a 12 Mbp large-scale chromosomal inversion distinguishing human and Sumatran orangutan chromosomes (see light blue alignment in Fig. 3a, b). In orangutan the corresponding breakpoint of synteny is anchored in one of the expanded *TBC1D3* clusters. This structure is syntenic with macaque, suggesting that it was the ancestral configuration, while human structure, shared with gorilla and chimpanzee, was derived. Similarly, one of the fission breakpoints of chromosome 17 resulting in gorilla chromosomes 4 and 19 (Stankiewicz et al. 2001) maps precisely to *TBC1D3* and *USP6* duplications in the gorilla lineage.

**Figure 3:**
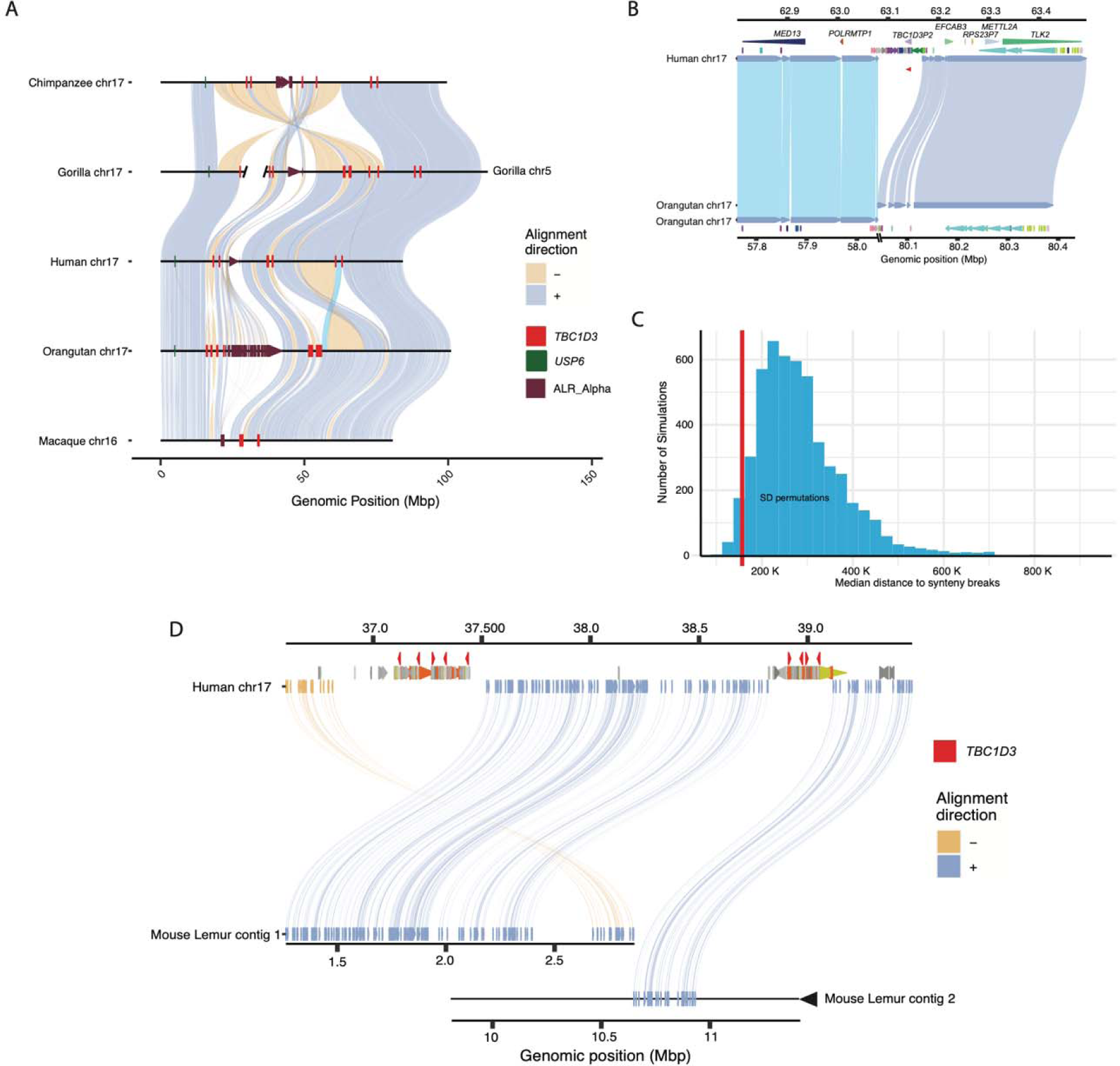
Large-scale chromosomal rearrangements and *TBC1D3* duplications. **(A)** Synteny plots of orthologous chromosome 17 in primates reveal syntenic blocks in direct (blue) and inverted (yellow) orientation. Alpha satellite sequence, *TBC1D3* copies, and *USP6*—a hominoid fusion gene of *TBC1D3*—are illustrated in maroon, red, and green, respectively. *TBC1D3* demarcates the boundaries of large-scale rearrangements on chromosome phylogenetic group XVII. **(B)** *TBC1D3* duplication block (cluster of colored arrows) demarcates the boundary of a 12 Mbp inversion between human and orangutan chromosomes. **(C)** Permutation test of segmental duplication proximity to synteny breaks. 5000 permutation tests were performed, where segmental duplication samples were taken and median proximity to breaks in synteny were measured. True *TBC1D3* mappings fall within the lowest 3% of permutations suggesting a nonrandom association between *TBC1D3* and breakpoints in synteny. **(D)** Synteny plot showing orthologous alignments between human *TBC1D3* and mouse lemur.

To test if the association with *TBC1D3* and breakpoints of synteny was significant, we developed a permutation test. We randomly selected an equivalent sequence and number of mappings throughout chromosome 17 for these five orthologous primate chromosomes and measured the median distance of these mappings to the nearest synteny break. In over 5000 permutation tests we never observed a distance as low as that of true *TBC1D3* mappings (Supplementary Fig. S8). We repeated the test by limiting our samplings to SD sites on chromosome 17. Even with this restriction, the observed distance to *TBC1D3* resided in the bottom 3% of the simulated distribution (Fig. 3c), suggesting a nonrandom association of *TBC1D3* SDs with large chromosomal rearrangements during primate evolution.

To assess the origin of *TBC1D3* gene clusters, we sequenced and assembled the genome of an outgroup primate species using HiFi data generated from a mouse lemur (*Microcebus murinus)* and identified two sequence contigs (2.8 Mbp and 14 Mbp) spanning the region (Fig. 3d). Both clusters 1 and 2 appeared to be absent; however, the corresponding regions demarcate breakpoints of synteny when compared to Old World monkey and ape lineages, suggesting *TBC1D3* is exclusive to the simian order.

### *TBC1D3* transcript and open reading frame (ORF) prediction

Gene model characterization of *TBC1D3* has been particularly challenging given the high sequence identity and variable nature of the duplicated genes. This has made it difficult to distinguish genes that are expressed and potentially functional from pseudogenes. To address this limitation, we sequenced high-fidelity, full-length non-chimeric (FLNC) cDNA using a PacBio isoform sequencing (Iso-Seq) assay (Dougherty et al. 2018; Methods). We generated or analyzed data from testis tissue of chimpanzee, gorilla, bonobo, Sumatran and Bornean orangutan (Makova et al. 2023) and from pooled human fetal brain tissue (Supplementary Table S3). Additionally, we analyzed a very deep pool of ∼500 million human FLNC recently generated from induced pluripotent stem cells (iPSCs) (Cheung et al. 2023). We mapped FLNC reads to both haplotypes of the respective species of origin genome assemblies, allowing only high-quality mappings, and tracking all best map assignments versus multiple mappings among the paralogous copies for each species (Fig. 4a; Methods). While unambiguous one-to-one assignments between transcripts and specific paralogs could not always be made, the analysis revealed three important features. First, *TBC1D3* is transcribed in all ape lineages with evidence of multiple paralogs expressed where there are duplications (Supplementary Fig. S4). Second, the canonical 14-exon gene model is retained across the apes, with evidence of exon exaptation and exon loss for a minority subset of transcripts in chimpanzee and Sumatran orangutan (Fig. 4a). Third, the predicted ORF is, in general, maintained. In humans, however, both transcription and ORF maintenance are most likely to be retained among *TBC1D3* copies mapping to clusters 1 and 2 in contrast to distal orphan copies (see Fig. 2a human chromosome 17 ideogram).

**Figure 4:**
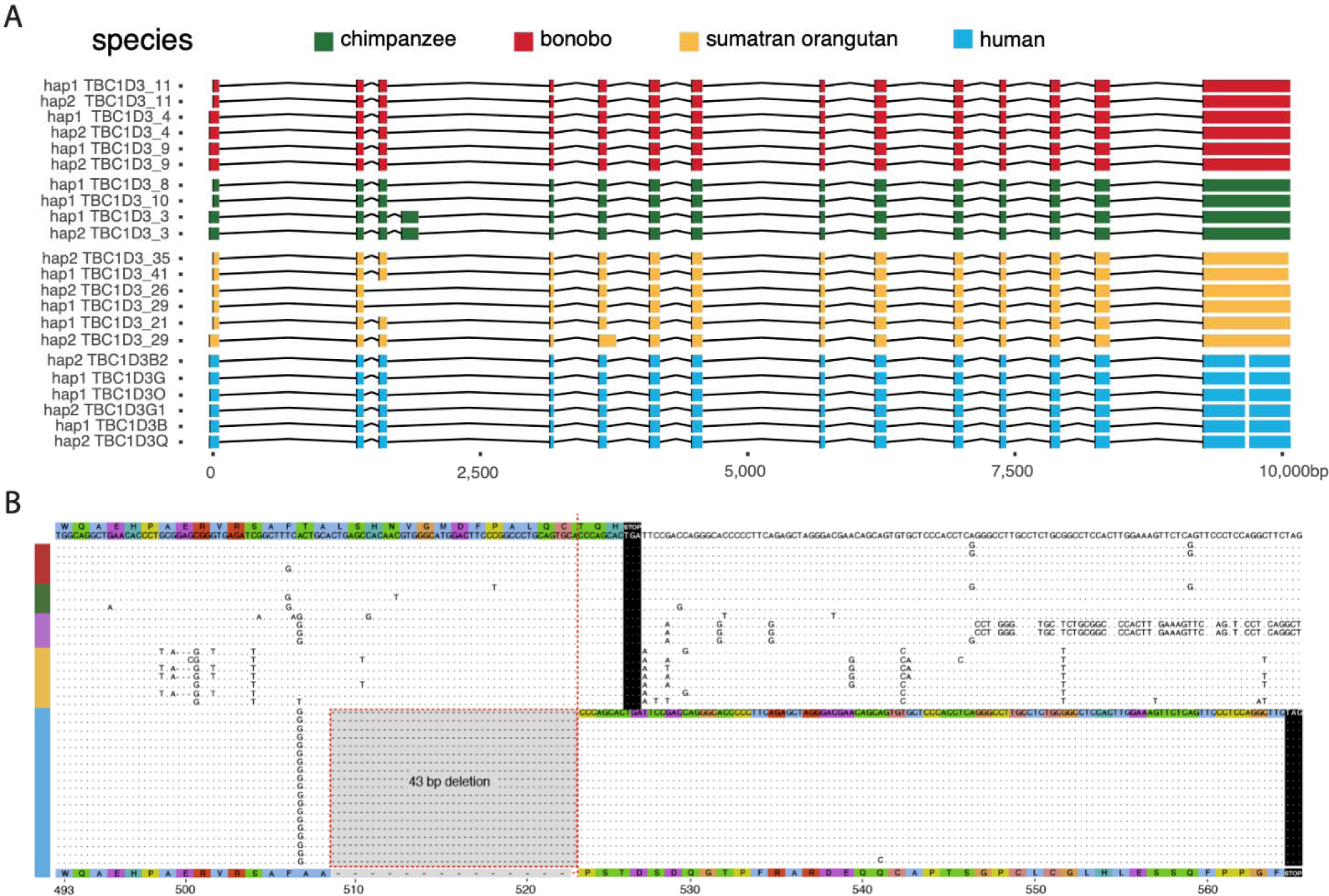
Human-specific C-terminus modification of *TBC1D3*. **(A)** The intron/exon structure of expressed *TBC1D3* isoforms with protein-encoding ORFs. Each row constitutes a paralog-specific isoform observed based on Iso-Seq (Methods). All isoforms were mapped to human *USP6* for a common reference. Exons are colored by species, with arches representing introns. **(B)** Amino acid sequence alignment of the carboxy terminus of expressed primate *TBC1D3* paralog sequences predicted from Iso-Seq full-length cDNA. All and only human-expressed copies contain a 43 bp deletion within the ORF of the terminal exon resulting in a frameshift, creating an extension of 41 novel amino acids to the C-terminus.

During our comparison of human and NHP *TBC1D3* gene models, we noted that all human transcripts harbor a 43 bp deletion in the ORF absent in other NHPs (Fig. 4b). This deletion removes the last 17 amino acid residues common to NHPs and introduces a frameshift resulting in a 41 amino acid extension and a novel carboxy terminus of human *TBC1D3*. All other NHPs lack this carboxy extension due to a shared common stop codon. We also confirmed this human-specific difference at the level of the assembly using prosplign (Kiryutin et al. 2017; Methods). Remarkably, the 43 bp deletion is restricted to all duplicate copies mapping to human *TBC1D3* clusters 1 and 2 and is not observed among the older orphan paralogs distributed throughout human chromosome 17 (Supplementary Fig. S5). These findings indicate that this fundamental change in the ORF is human specific and occurred during human *TBC1D3* expansion within clusters 1 and 2. We predicted the effect of this modification on the tertiary structure of *TBC1D3* using Alphafold2 but found that the novel C-terminus sequence was disordered (Supplementary Fig. S5; Jumper et al. 2021).

### African ape positive selection

Using the full-length transcript isoforms that were generated and mapped to the complete genome assemblies from each primate (Fig. 5a), we constructed MSAs of both the intronic regions and codon-aligned exonic regions (Methods). We tested for a significant excess of amino acid replacements with a maximum likelihood branch-model test (Fig. 5b; Yang 2007) and found strong statistical support for positive selection among ancestral branches leading to African ape lineages. This positive selection is detected only for *TBC1D3* copies mapping to clusters 1 and 2 and not among orphan copies or other ape clusters distributed along chromosome 17. Specifically, positive selection was detected along three sequential branches, starting from a branch ancestral to all African ape cluster 1 and 2 *TBC1D3* paralogs (marked in red), and terminating along the human-specific clade (p-value = 0.024). This selection appears to have occurred after divergence from orangutan, and after an African ape-specific translocation of *TBC1D3* paralogs to chromosome 17q23 (Fig. 3a). Neither orangutan copies expressed from clusters 1 and 2, nor expressed chimpanzee/bonobo copies mapping distally to clusters 3 and 4 (yellow), show signatures of positive selection. Focusing on African ape copies mapping to clusters 1 and 2, we tested for site-specific signatures of positive selection on amino acid residues. Using a Bayesian posterior probability cutoff of 0.8, we identified six sites of positive selection, with the strongest signals mapping within the TBC/Rab GTPase-activating protein (GAP) domain, as well as two residues proximal to the carboxy terminus of *TBC1D3* (Fig. 5c).

**Figure 5:**
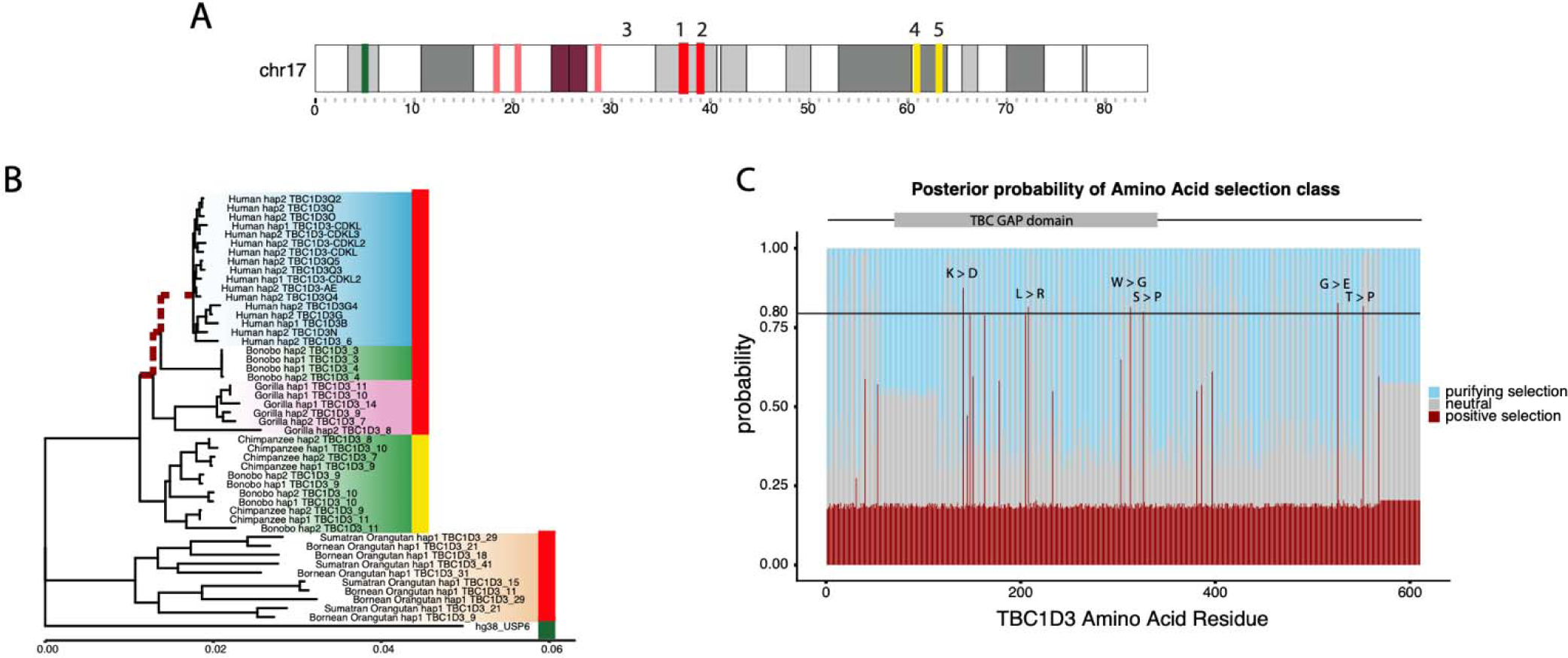
Positive selection of the *TBC1D3* gene family. **(A)** Chromosome 17 ideogram marking *TBC1D3* expansion clusters and distal loci expressed in chimpanzee and bonobo. **(B)** Branch model test of selection for expressed *TBC1D3* paralogs. Neutral, maximum likelihood phylogeny corresponding to the introns of expressed *TBC1D3* paralogs used as a baseline to compute branch model test of selection in coding sequence. Red dashed branches represent branches identified under positive selection (p-value = 0.024). Colored bars on right of phylogeny illustrate location of origin of *TBC1D3* copies in (A), red indicating paralogs from clusters 1 and 2, yellow marking expressed paralogs from distal q-arm expansions 3 and 4, and green marking *USP6*, the outgroup. **(C)** Branch site model test across *TBC1D3*. A branch site model was conducted using the codon alignment of the same *TBC1D3* expressed isoforms, with the three ancestral branches leading to gorilla, chimpanzee, bonobo, and human *TBC1D3* copies as the foreground, and all other branches as the background. Posterior probabilities for positive, neutral, and purifying selection are illustrated in red, gray, and blue, respectively, with red indicating sites under selection in the foreground branches (omega = 35).

### Pangenomic characterization and transcription of human *TBC1D3* copies

Given the extraordinary copy number variation among human copies mapping to clusters 1 and 2, we applied a pangenomic approach to organize and characterize human paralogs. We initially constructed pangenome graphs with Minigraph from the sequence-resolved human haplotypes. However, few paralogs were grouped as common or shared but instead the majority of *TBC1D3* copies were represented as isolated nodes with single-haplotype support (Supplementary Fig. S3; Li et al. 2020). As a result, we applied a phylogenetic approach that organized *TBC1D3* copies into groups where genetic distance exceeded the expected level of intra-allelic variation (Methods). We defined 11 distinct phylogenetic groups (Fig. 6a) and named them based on *TBC1D3* paralogs already present in the human reference genome (GRCh38; Supplementary Fig. S7). In some cases, multiple distinct paralogs were placed into the same phylogenetic group if paralogous variation was less than the expected extent of allelic variation (e.g. *TBC1D3-AE* or *TBC1D3-CDKL*). We identified four novel phylogenetic groups representing paralogous copies not present in the human reference genome assembly: *TBC1D3M, TBC1D3N, TBC1D3O,* and *TBC1D3Q.* Most phylogenetic groups are distributed across human continental population groups and are specific to either cluster 1 or 2. *TBC1D3F*, however, is exclusive to Amerindians and maps to cluster 2 yet has greater homology to cluster 1 *TBC1D3* members. Interestingly, a detailed examination of the genomic organization of one of these Amerindian haplotypes, HG01109 H2, reveals that the entire 1.35 Mbp region bracketed by clusters 1 and 2 has been inverted, suggesting that inversion, as well as gene conversion, may be playing a role in relocating *TBC1D3* paralogs between clusters 1 and 2 (Fig. 6b).

**Figure 6:**
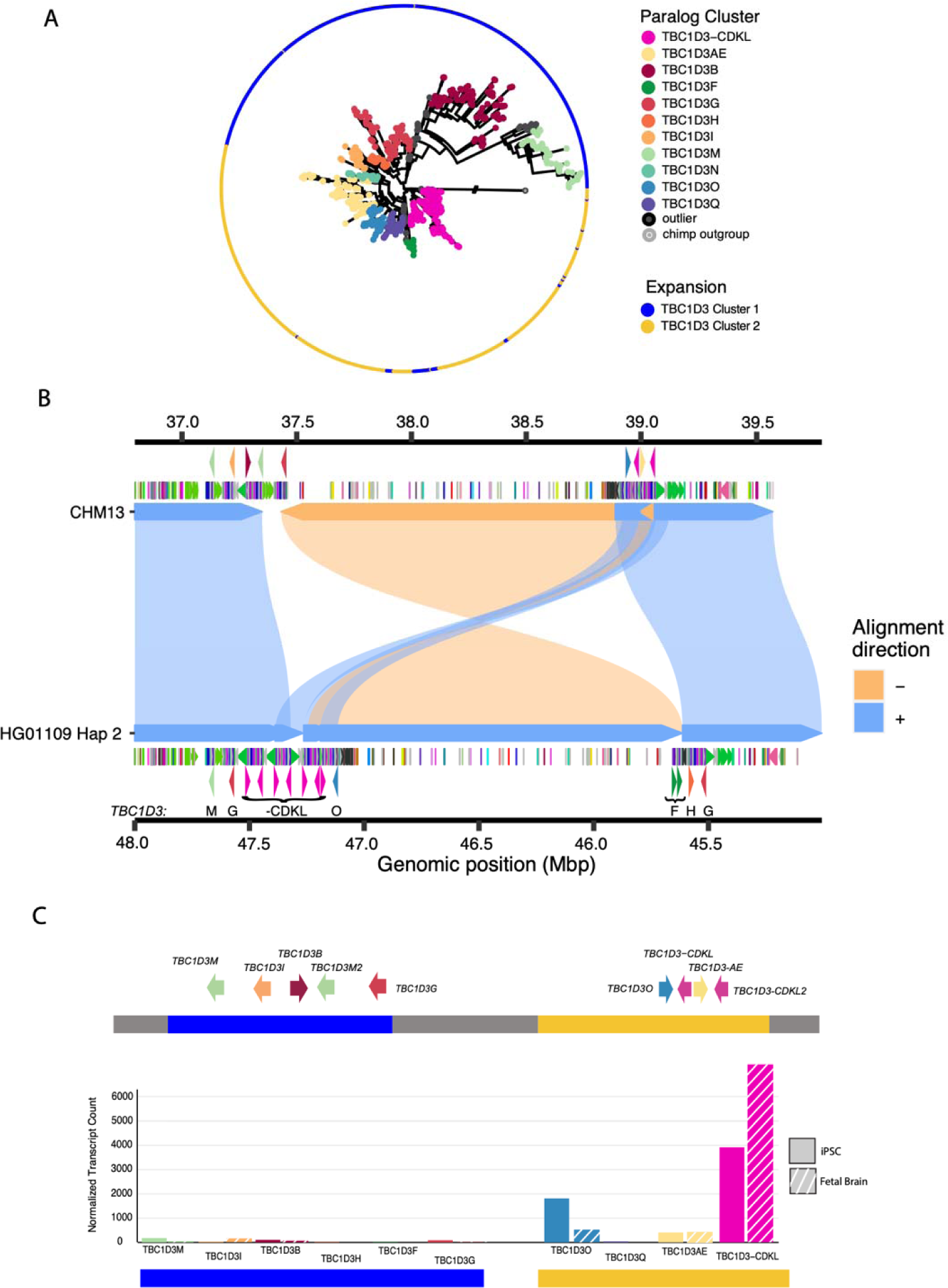
Pangenomic characterization and expression of *TBC1D3* in humans. **(A)** Maximum likelihood phylogeny of all validated *TBC1D3* cluster 1 and 2 paralogs in humans, outgrouped to chimpanzee *TBC1D3*. Individual cluster paralogs were identified by limiting intra-cluster variation to 1.5× allelic variation observed in SD sequence. This resulted in a gene family of 11 common paralogs. **(B)** Inversion haplotype of HG01109 hap2 (bottom) aligned to CHM13 (top). **(C)** Visual illustration of CHM13 clusters 1 and 2 with new paralog characterization, and expression of these paralogs across iPSCs and fetal brain Iso-Seq libraries, normalized to median haplotype paralog copy number.

Using this phylogenetic group classification of cluster 1 and 2 members, we revisited expression of the *TBC1D3* gene family in humans, taking advantage of the deep Iso-Seq datasets that had been generated from both iPSCs and fetal brain (Supplementary Table S3). We mapped FLNC reads from both sources to the phylogenetic pangenome groups and identified the best primary paralog mapping for each read (Methods). We find that the majority of *TBC1D3* expression— 91% in iPSCs and 96% in fetal brain—originates from paralogs assigned to cluster 2 specifically. Furthermore, the majority of this sequence—89% in fetal brain and 69% in iPSCs—maps to a single phylogenetic group: *TBC1D3-CDKL*. This enriched paralog expression is consistent, even when normalized by median *TBC1D3* paralog copy (Fig. 6c). This paralog expression exclusivity may explain why a gene family predicted to be critical to cortical expansion may be so variable in copy number and structure among humans.

## DISCUSSION

Long-read sequencing and advances in *de novo* genome assembly have enabled comprehensive characterization of complex, duplicated loci (Liao et al. 2023). Here, we investigated the evolution and transcription of *TBC1D3*, a “hominoid-specific” gene family functionally implicated in the proliferation of neuronal progenitors and cortical expansion and folding of the human brain (Ju et al. 2016; Paulding et al. 2003; Sudmant et al. 2010; Hou et al. 2021). Previous analyses based on short-read sequencing read depth revealed that *TBC1D3* was highly copy number polymorphic; however, its genome organization was never fully realized in large part due to high sequence identity of the different paralogs (Sudmant et al. 2010). Using Hifiasm and Verkko, we successfully assembled and validated 69 human haplotypes from three references and 33 human samples across *TBC1D3* clusters 1 and 2 (Cheng et al. 2021; Rautiainen et al. 2023). While both methods combine long-read sequencing data to traverse these large gene-rich repeat regions, neither method fully resolved all structures. Use of Verkko following application of Hifiasm did, however, recover an additional ∼30% (20/66) of *TBC1D3* genome structures (Fig. 1a) in large part due to the incorporation of ultra-long ONT sequence data. We find that the human *TBC1D3* gene family is among the most copy number variable gene families, with over 60% of human haplotypes containing a unique structural configuration at each cluster with an overall structural heterozygosity estimated at 90%. *TBC1D3* copy number at each cluster ranges from 1 to 14, with samples from African populations tending to have an overall higher copy number when compared to non-Africans. This observation is consistent with previous investigations of increased SD copy number in African haplotypes (Vollger et al. 2023). While graph-based approaches such as Minigraph failed to produce a biologically meaningful representation of this locus, a phylogenetic analysis identified 11 common *TBC1D3* groups—four of which were novel and not represented in either the GRCh38 or CHM13 T2T human references (Fig. 6a; Supplementary Fig. S3).

Given the extreme structural diversity of this locus, how could a gene implicated in a process as critical as brain cortical expansion during human evolution be so variable within the human population? Leveraging a deep long-read Iso-Seq resource, which allowed the paralog isoforms to be distinguished, we investigated the relative transcription of the *TBC1D3* paralog groups in two developmental contexts (iPSCs and fetal brain). We find that *TBC1D3* paralogs mapping to cluster 2, most notably *TBC1D3-CDKL*, account for ∼90% of assigned transcripts. We hypothesize that this restricted pattern of expression may explain how such high *TBC1D3* copy number variation is tolerated, because only one or two copies, located at the telomeric end of *TBC1D3* cluster 2, are exclusively expressed. This model of regulation is reminiscent of the green opsin gene family on the X chromosome, where a single locus control region promotes expression of the most proximal green opsin paralog, while downstream duplicates are transcriptionally silent (Hayashi et al. 1999). In this model, many of the other *TBC1D3* paralogs are either inactive pseudogenes or “genes-in-waiting”. Notably, for 67 of the 69 assembled haplotypes, this expressed *TBC1D3* paralog is the last copy in cluster 2 and oriented such that the unique sequence flanking the telomeric end of the cluster is directly upstream to its transcription start site. A genome-wide analysis identified that the 20 kbp of flanking unique sequence immediately telomeric of this paralog group falls within the lower 5% for pairwise nucleotide diversity and may reflect either a selective sweep or regulatory sequence under strong purifying selection (Supplementary Table S9).

*TBC1D3* is just one example of approximately two dozen core duplicons—originally defined as sequence overrepresented in SD repeat graphs, marking the focal point of interspersed duplication blocks (Jiang et al. 2007, Marques-Bonet & Eichler 2009; Dennis et al. 2017). Several core duplicons have been associated with recurrent and independent duplications in primates, chromosomal rearrangements among apes, large-scale inversion polymorphisms in humans, and developmental disorders (Johnson et al. 2006; Mao et al. 2024; Nuttle et al. 2016; Zody et al. 2006; Mohajeri et al. 2016; Porubsky et al. 2022; Maggiolini et al. 2019; Antonacci et al. 2010). *TBC1D3* is no exception. Our data suggest lineage-specific duplication and expansion in different primate lineages (Fig. 3). Thus, the expansion of this gene family is not hominoid specific, being clearly present in the Old World monkey genomes. *TBC1D3* clusters 1 and 2 demarcate the boundary of recurrent microdeletions associated with renal cyst and diabetes syndrome (RCAD) on chromosome 17q12 (Mefford et al. 2007). The clusters also contain the breakpoints of a 2.2 Mbp inversion polymorphism that we discovered in this study in one Amerindian haplotype, consistent with ongoing nonallelic homologous recombination between inverted *TBC1D3* gene clusters. Similar to many of the genes embedded within core duplicons, such as *NPIP*, *RGPD*, *NBPF*, among others, we present evidence of positive selection in African ape *TBC1D3* based on dN/dS (Johnson et al., 2006; Mao et al., 2024; Popesco, 2006; Bekpen 2019). We also show human-exclusive modification of the carboxy terminus. This signature is potentially consistent with recent neofunctionalization and diversification of this gene family in humans. It is interesting that a recent analysis of the putative protein structure encoded by 10 of the core duplicons, including *TBC1D3*, are enriched for disordered regions that may enhance protein-protein interactions or post-translational modification (Bibber et al. 2020).

In this study, we estimate that *TBC1D3* expanded ∼2.5 mya in humans—a time point according to the fossil record when the genus *Homo* transitioned from *Australopithecus*, preceding the onset of frontal cortical expansions in *Homo habilis* (Spoor et al. 2015). Functional investigations have suggested different biochemical roles for TBC1D3 at the cellular level, all of which increase cell proliferation. Two functions occur in the cytosol, where TBC1D3 antagonizes ubiquitination and degradation of EGFR and INS-1 receptors, driving cell proliferation in cell culture (Wainszelbaum et al. 2008; 2012). The third, in contrast, proposes that TBC1D3 is shuttled to the nucleus in neuron progenitor cells where it antagonizes *G9a* methyltransferase and, as a result, epigenetically inhibits neural progenitor differentiation (Hou et al. 2021). We propose that the human-specific modified carboxy terminus, as well as the positively selected amino acid changes, play a critical role in these adaptive functions by potentially directing novel post-translational modifications or altering the localization and trafficking of TBC1D3 proteins (Sharma and Schiller 2019). It will be important to compare the protein structure and function of human and NHP *TBC1D3* paralogs to determine if neofunctionalization has indeed occurred as a result of these changes in the human lineage. Finally, it will also be important to understand how *TBC1D3* clusters are transcriptionally regulated in relevant cell types during development, given the extraordinary copy number variation present in the human population. Our results suggest a strong position effect from the most telomeric copies located in cluster 2 and long-read sequencing-based methods, such as Fiberseq, may be particularly useful to dissect the regulatory architecture of the clusters (Stergachis et al. 2020). We hypothesize that most paralogs are transcriptionally repressed or expressed at a much lower level. It is possible that residual expression of other *TBC1D3* paralogs in each individual may subtly modify expression and development in humans. The long-read genomic resource we developed provides the framework to address these and other functional questions related to the emergence of this intriguing gene family.

## METHODS

### Long-read sequence and assembly

The majority of genomes used in this study were sequenced previously as part of other assembly efforts to generate phased genomes or T2T genomes and are publicly available (Liao et al. 2023; Mao et al. 2024; Makova et al. 2023). See Supplementary Table S2 for species, coverage, and project details. This study focused only on analyzing sequence contigs that contained copies of *TBC1D3* paralogs, and we evaluated each contig for gaps and contiguity (see below). Most human genomes were originally assembled using Hifiasm (version 0.15.2), but *TBC1D3-* containing contigs that failed QC were reassembled with Verkko (versions 1.0, 1.1, 1.2, and 1.4) using a combination of both HiFi and ONT sequence. In general, haplotypes were phased using parental k-mer information where available, or Hi-C chromatin capture data (Auton et al. 2015; Kronenburg et al. 2021). For the chromosome 17 comparison, it was observed that the macaque orthologous chromosome was fragmented and was subsequently scaffolded using RagTag (version 2.1.0) with the Mmul10 reference as the scaffold (Alonge et al. 2022; Hughes et al. 2012). In this study, we generated assemblies for only two species: gelada (*Theropithecus gelada*) and mouse lemur (*Microcebus murinus*). High molecular weight DNA was prepared from peripheral blood of a male gelada (*DRT_2020_14_TGE*) and from skin fibroblasts of a female mouse lemur (*Inina_MMUR*). HiFi sequence data (50×, 30×) were generated using the Sequel II platform and assemblies were generated with Hifiasm (Supplementary Table S2).

### Assembly validation

#### Illumina copy number validation

Sample assemblies were first validated using diploid assembly *TBC1D3* copy number estimates to Illumina sequence copy number estimates, an orthogonal sequencing approach (Supplementary Fig. S1). Sample genome haplotypes were merged and k-merized into 32 bp k-mers using Meryl (version 1.3). In parallel, sample Illumina sequence libraries were similarly k-merized into 32 bp with Meryl. Next, k-mer libraries were aligned to the T2T reference genome using FastCN, allowing for up to two mismatches between k-mer and assembly alignments (Nurk et al. 2022; Pendleton et al. 2018). We estimated the copy number of *TBC1D3* by taking the average copy number over one *TBC1D3* paralog, *TBC1D3L*, and compared these estimates against one another in a scatter plot (Fig. 1a, Supplementary Fig. S1).

#### Self-read mapping validation

We also applied NucFreq (Vollger et al. 2018) to assess the integrity of each *TBC1D3* assembly. Each sample’s respective HiFi sequencing libraries were trio phased using Canu (version 2.1.1; Koren et al. 2017) and mapped back onto their respective *de novo* assemblies. To qualitatively validate assembly, we plotted sequence depth of both the primary and secondary base of reads aligned over the *TBC1D3* expansions (Supplementary Fig. S9). First, we removed samples with obvious gaps over the *TBC1D3* expansion 1 and 2 loci, which could be identified if the locus was broken across multiple contigs or if the assemblies had a lack of HiFi sequence support over a given region. Next, we identified assemblies with collapses over the *TBC1D3* expansion 1 and 2 regions by looking at secondary base read depth. HiFi sequencing is 99.9% accurate, with occasional low-frequency false base calls. Our expectation is that this frequency can be observed over a given region as the secondary base, remaining well below 1% frequency. Any haplotypes with a noticeable increase in secondary base frequency over particular stretches were marked as collapsed. Usually, these samples included a spike in primary base coverage as well as over the collapsed region. Additionally, Hifiasm samples were validated with GAVISUNK (Dishuck et al. 2022). Phased ONT reads were mapped over each sample’s respective assemblies and singly unique nucleotide k-mer anchors were marked. We expect, for correct assemblies, that every region of the assembly will be supported by at least one ONT sequence, which is not used during Hifiasm assembly. Any locations with a gap in ONT assemblies were marked as not validated.

### Repeat and gene mapping annotation

We defined repeat content in the genome using TRF (version 4.09) for simple tandem repeats, RepeatMasker (version 4.1.2-p1) for common transposon and retrotransposon elements, and DupMasker to define duplicons associated with human SDs (Jiang et al. 2008). *TBC1D3* loci were identified in the GRCh38 reference genome based on RefSeq annotations and mapped to other assemblies using Minimap2 (version 2.24), using the asm20 standardized setting and allowing for up to 1000 secondary alignments (Li et al. 2018). These mappings were filtered to contain at least 6 kbp of sequence, over half the length of the canonical *TBC1D3* gene model. For more distantly related lineages, including the New World monkeys, we mapped *TBC1D3* sequence using blat (version 3.5), allowing a maximum intron length of 5 kbp, half the *TBC1D3* gene model length, and a minscore of 100. These relatively loose mapping constraints identified many candidate *TBC1D3* paralogs, more than expected by either Illumina- or assembly-based *TBC1D3* copy number estimates, that were subsequently filtered based on expression, divergence, or minimum length match.

### Structural variation and heterozygosity characterization

Validated cluster 1 and 2 *TBC1D3* haplotypes were aligned to one another in an all-by-all fashion using Minimap2 (version 2.24) auto settings -x asm5, allowing up to 1 kbp of insertions in cigar strings. We labeled two haplotypes as structurally equivalent if 90% or more of their sequence could be mapped to one another in a single alignment. We repeated this exercise for all pairs of haplotypes, calculated the number of valid haplotypes with no structurally equivalent pair, and divided by the total number of validated haplotypes to determine our structural variation statistic. For structural heterozygosity we identified all samples whose two haplotypes were not structurally equivalent and divided by the total assembled samples. Contig and chromosome alignments (e.g., Figs. 3 & 5) were visualized by SVByEye using either plotMiro for pairwise alignment, or plotAVA for all-vs-all alignments (https://github.com/daewoooo/SVbyEye). Blue alignments represent directly orientated alignments, and yellow indicates inverted alignments. For local *TBC1D3* structure comparison (Supplementary Fig. S6), we extracted primate *TBC1D3* copies, along with 25 kbp of flanking sequence, from five primate lineages and mapped to one another. These copies were organized to reflect the closest alignments, both by length and identity.

### *TBC1D3* breakpoint simulation

We mapped orthologous chromosome 17 relationships and annotated *TBC1D3* copies using Minimap2 -x asm20. Synteny was annotated using Asynt get.synteny.blocks.multi command, with max_gap = 200000, min_block_size = 1000000, and min_subblock_size = 50000, producing a tab-delimited file marking the target and query breaks of blocks. For each *TBC1D3* copy we identified the nearest synteny break along the respective chromosome, then computed median distance to synteny breaks of all *TBC1D3* mappings. Next, we conducted a permutation experiment. For each primate orthologous chromosome 17, we randomly selected ∼11 kbp blocks at the same quantity as the number of *TBC1D3* mappings observed in the respective primate chromosome. We repeated the median distance experiment and plotted the distribution of 5000 permutations.

### Multiple sequence alignment (MSA)

Sequence was extracted from assemblies by mapping *TBC1D3* sequence to full genome assemblies with Minimap2 (version 2.24) and extracting the mapped reference sequence with BEDTools (version 2.29.2) (Li 2018; Quinlan and Hall 2010). MSAs were constructed with MAFFT with parameters --reorder --maxiterate 1000 --thread 16 (version 7.453; Katoh et al. 2002). Following MSA construction, spurious alignments were pruned with trimmal (--gappyout) (version 1.4) and manually trimmed. Codon alignments were generated with matched ORF and amino acid sequence fasta files. First, an amino acid MSA was generated with MAFFT. Then, the ORF fasta was aligned to the amino acid MSA with pal2nal (Suyama et al. 2006).

### Phylogenetic analyses

Maximum likelihood phylogenies were generated with iqtree2 using model setting -m MFP, 1000 lrt replicates, and -b 1000 replicates for bootstrap (version 2.1.2). Additionally, each phylogeny generated was outgrouped to a sequence: marmoset *TBC1D3* for primate phylogenetic analysis, and chimpanzee *TBC1D3* for human paralogs clustering. Phylogenetic trees were illustrated in R with ggtree (Yu 2022). Timing estimates for individual primate expansions were conducted using BEAUTi for data input and BEAST2 for computation (Drummond et al. 2012; Bouckaert et al. 2019). We used human–macaque and human–chimpanzee divergence times of 25 and 6.5 mya, estimated by the fossil record, as benchmarks for the computation (Stevens et al. 2013; Dunsworth et al. 2010). With these references, we calculated the 95% confidence intervals of mutation rate within sequences, then estimated species-specific expansions with this mutation rate as well as branch lengths of the primate phylogeny. For tests of positive selection, we isolated intronic sequence and exonic sequence from paralog isoforms with expression support from human, chimpanzee, bonobo, gorilla, Sumatran orangutan, and Bornean orangutan genome assemblies. Next, we generated an intronic MSA and exonic codon aligned MSA as described above. With the intronic MSA, we generated a maximum likelihood phylogeny outgrouped to human *USP6*. With intronic phylogeny and codon-aligned MSA, we tested all branches for selection in a free-ratios test. We ignored predicted dN/dS values for terminal branches, as they occurred too recently to effectively test for selection. Among deeper branches we identified three that were putatively under selection, as mentioned in the results. We tested these for significance using a likelihood ratio test in R. Next, we isolated these branches in a branch-site model test, setting these three branches as the foreground, and all other branches as background. We selected amino acid residues under selection based on the Bayes Empirical Bayes posterior probability.

### Iso-Seq and transcript analyses

Primate Iso-Seq testis data was generated by Makova et al. 2023. Similarly, human iPSC Iso-Seq was previously generated by Cheung et al. 2023. Fetal brain tissue was derived from 59 spontaneously aborted fetuses. This sequence was enriched for both *TBC1D3* and *NPIP*, using the hybridization capture protocol described in Dougherty et al. 2018, with probes provided in Supplementary Table S10. FLNC libraries were mapped to respective species libraries with Minimap2 using parameters -ax splice --sam-hit-only --secondary=yes -p 0.5 --eqx -K 2G -G 8k -N 20. FLNC libraries were first filtered for reads greater than or equal to 1000 bp in length and with sequence quality of 99.9% or greater. Each library was subsequently mapped to the genome assembly corresponding to the respective species of origin using SAMtools and BEDTools. Next, we determined which *TBC1D3* paralogs were likely expressed by selecting paralogs with read support with mapping quality greater than or equal to 99.9% sequence identify. These reads were subsequently reduced into common isoforms with isoseq3 collapse, and ORFs were predicted with Orfipy (Singh and Wurtele 2021). For primate *TBC1D3* gene model comparison, isoforms with at least three independent reads of support and with the longest maintained ORF were compared. We required these reading frames to span within 100 bp of the canonical *TBC1D3* start and stop as defined by RefSeq (O’Leary et al. 2016). Human FLNC reads from fetal brain and iPSCs were mapped to all validated human haplotypes. Next, we compared these primary alignments to one another and considered the cluster paralog from which they were derived. Any Iso-Seq read with primary minimap2 alignment scores of 10 or greater for a given paralog cluster relative to all other cluster mappings were retained, while other mappings were marked as ambiguous and ignored.

### Analysis of coding sequence

To validate the observed deletion of coding sequence in humans, we selected human TBC1D3L amino acid sequence and mapped this sequence to all genome assemblies with prosplign (Kiryutin et al. 2017). Prosplign is a tool that predicts DNA sequence representing the codons for a given protein amino acid sequence. This tool predicts splice junctions, as well as start and stop codons, and illustrates AA substitutions, frameshift mutations, and deletions in the underlying nucleotide sequence that is inconsistent with the provided amino acid sequence. We predicted human *TBC1D3* tertiary structure using the EMBL-EBI Alphafold2 database (Jumper et al. 2021; https://alphafold.ebi.ac.uk/). The predicted tertiary structure was illustrated using PyMol.

### Human pangenome graph construction

We built a pangenome graph of *TBC1D3* with Minigraph (version 0.20; Li et al. 2020), with settings -S -xggs -L 250 -r 100000 -t 16. We attempted graph construction with lower -l and -g settings as well but consistently observed that most haplotype *TBC1D3* paralogs were isolated to nodes without any allelic overlap from other human haplotypes.

### Human *TBC1D3* paralog grouping

We generated a phylogeny with the whole *TBC1D3* sequence for all cluster 1 and 2 copies identified in validated human assemblies, outgrouped to chimpanzee *TBC1D3*. We defined a heuristic cutoff based on allelic variation to define our clusters. Vollger et al. previously predicted allelic variation of 15.3 single-nucleotide variants per 10 kbp. We recursively identified clades with intra-variation of up to 1.5 times the allelic variation identified in SDs (Vollger et al. 2023). Additionally, we required that a given cluster have at least 10 independent paralogs of representation to be defined as a population-level paralog group.

## DATA ACCESS

Human Pangenome Reference Consortium samples were previously published, and sequencing and genome data are available on AnVIL (https://anvilproject.org) in the AnVIL_HPRC workspace. HG02723 was sequenced as part of the UCSC Genome Diversity Panel, available on their github website: https://github.com/human-pangenomics/hpgp-data. Primate T2T genome assemblies were previously published and made available on GenomeArk: https://genomeark.s3.amazonaws.com/index.html?prefix=species/. Gorilla Iso-Seq data is available as part of NCBI BioProject PRJNA902025 and in the Sequence Read Archive (SRA) database with identifier SRX18421140. Iso-Seq data is available in SRA for the following: bonobo: SRX18280098; chimpanzee: SRX18280097; Sumatran orangutan: SRX19199753 and SRX18421142; Bornean orangutan: SRX18421141. Sequencing and assembly data are available in NCBI under the following BioProject names: owl monkey: PRJNA941350; macaque: PRJNA877605; marmoset: PRJNA941358; gorilla: PRJNA916732 and PRJNA916733; bonobo: PRJNA916735 and PRJNA916734; chimpanzee: PRJNA916736 and PRJNA916737. Human iPSC Iso-Seq libraries were generated in Cheung et al. 2023 and made available in the dbGAP database under accession code phs002206.v4.p1. Fetal brain tissue sequenced is from BioSample SAMN09459150, and sequencing data made available under SRA SUB14282898. Gelada sequence and assembly data are available as part of NCBI BioProjects PRJNA1081468 and PRJNA1081469. Sequence data of gelada is available under SRA SUB14276633. Mouse lemur sequence and assembly data are available as part of NCBI BioProjects PRJNA1082315 and PRJNA1082316. Sequence data of the mouse lemur assembly are available under SRA SUB14282870.

## ACKNOWLEDGMENTS

We thank Noah Snyder-Mackler, Kenny Chou, and the Simien Mountains Gelada Research Project for providing peripheral blood for the sequence and assembly of the *Theropithecus gelada* genome. We thank Mark Krasnow for access to material from the mouse lemur (*Microcebus murinus).* We thank the Primate T2T Consortium, especially Kateryna Makova and Adam Phillippy, for providing us with early access to the high-quality ape genome assemblies. We thank Tonia Brown, Zoe Poyen, and Gerta Janss for manuscript proofreading and editing. We thank generous donors to Children’s Mercy Kansas City and Genomic Answers for Kids program supporting the human iPSC Iso-Seq (TP).

This article is subject to HHMI’s Open Access to Publications policy. HHMI lab heads have previously granted a nonexclusive CC BY 4.0 license to the public and a sublicensable license to HHMI in their research articles. Pursuant to those licenses, the author-accepted manuscript of this article can be made freely available under a CC BY 4.0 license immediately upon publication.

## Author Contributions

X.G. and E.E.E. conceived the project; X.G. assembled genomes, performed QC analyses, and conducted the analyses relevant to all manuscript figures; D.P. identified the HG01109 human inversion; D.Y. computed nucleotide diversity of unique sequence flanking *TBC1D3* clusters; M.L.D. conducted the probe capture experiments for fetal brain Iso-Seq data; P.C.D. provided technical and scientific consultation and assisted in Iso-Seq library processing; K.M.M. and A.P.L. generated PacBio HiFi and Iso-Seq sequencing data; K.H. and J.K. generated ONT sequencing data; S.C. processed samples and DNA necessary for mouse lemur genome assembly; T.P. generated iPSC Iso-Seq libraries; X.G. and E.E.E drafted the manuscript. All authors read and approved the final manuscript.

## SUPPLEMENTARY FIGURES

**Supplementary Figure S1:**
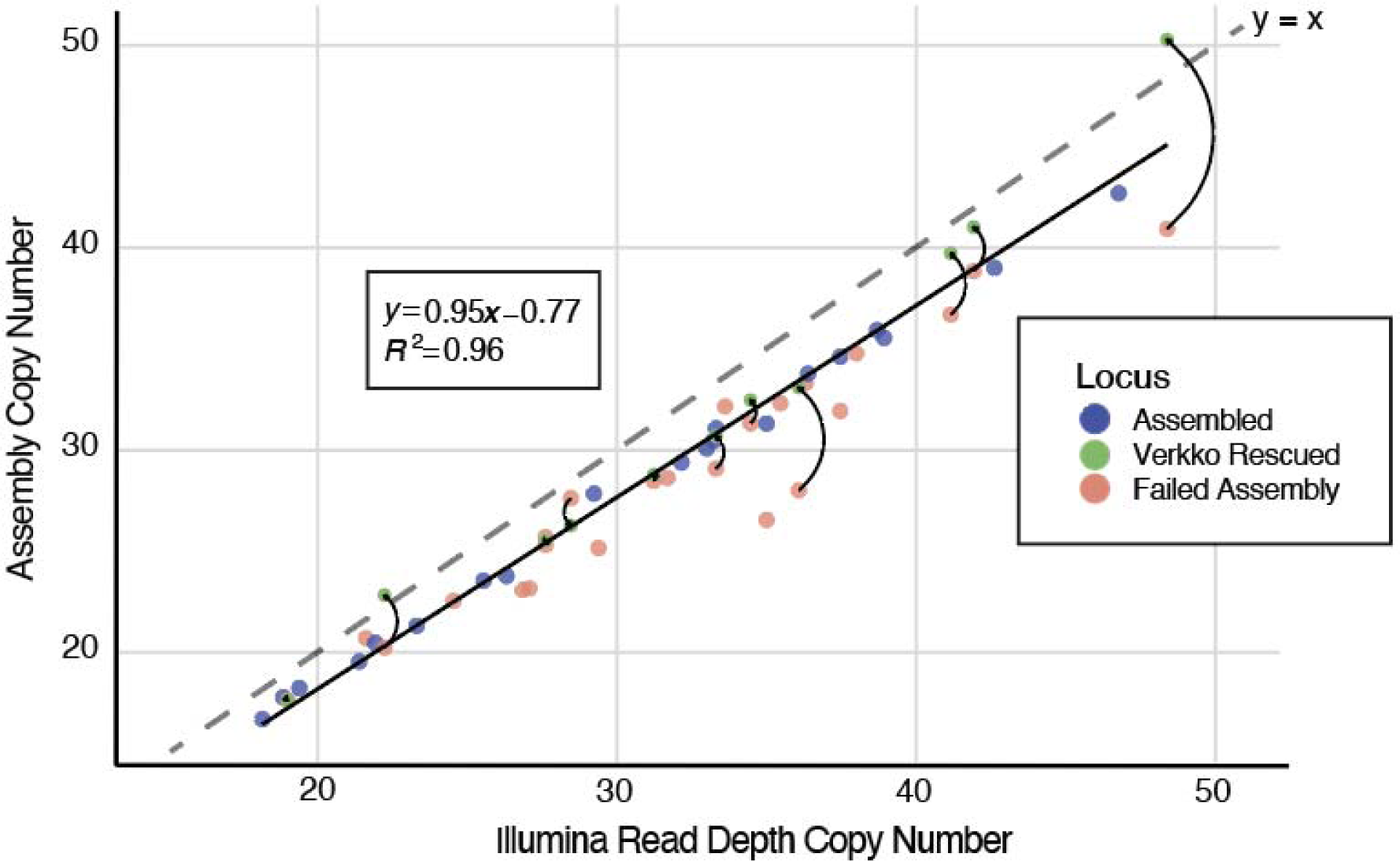
Assembly and rescue of *TBC1D3* phased haplotypes. Samples were first phased and assembled using exclusively HiFi sequence. The y-axis represents the diploid copy number of these assemblies, and the x-axis represents diploid copy number based on Illumina sequence, with y=x axis marked with a dashed line. Assemblies were validated by read depth estimates of HiFi and ONT, colored in blue if validated, and salmon colored if failing validation. We attempted to rescue sample haplotypes using a novel assembly approach leveraging both HiFi and ultra-long ONT. These samples, assembled by Verkko, are indicated in green.

**Supplementary Figure S2:**
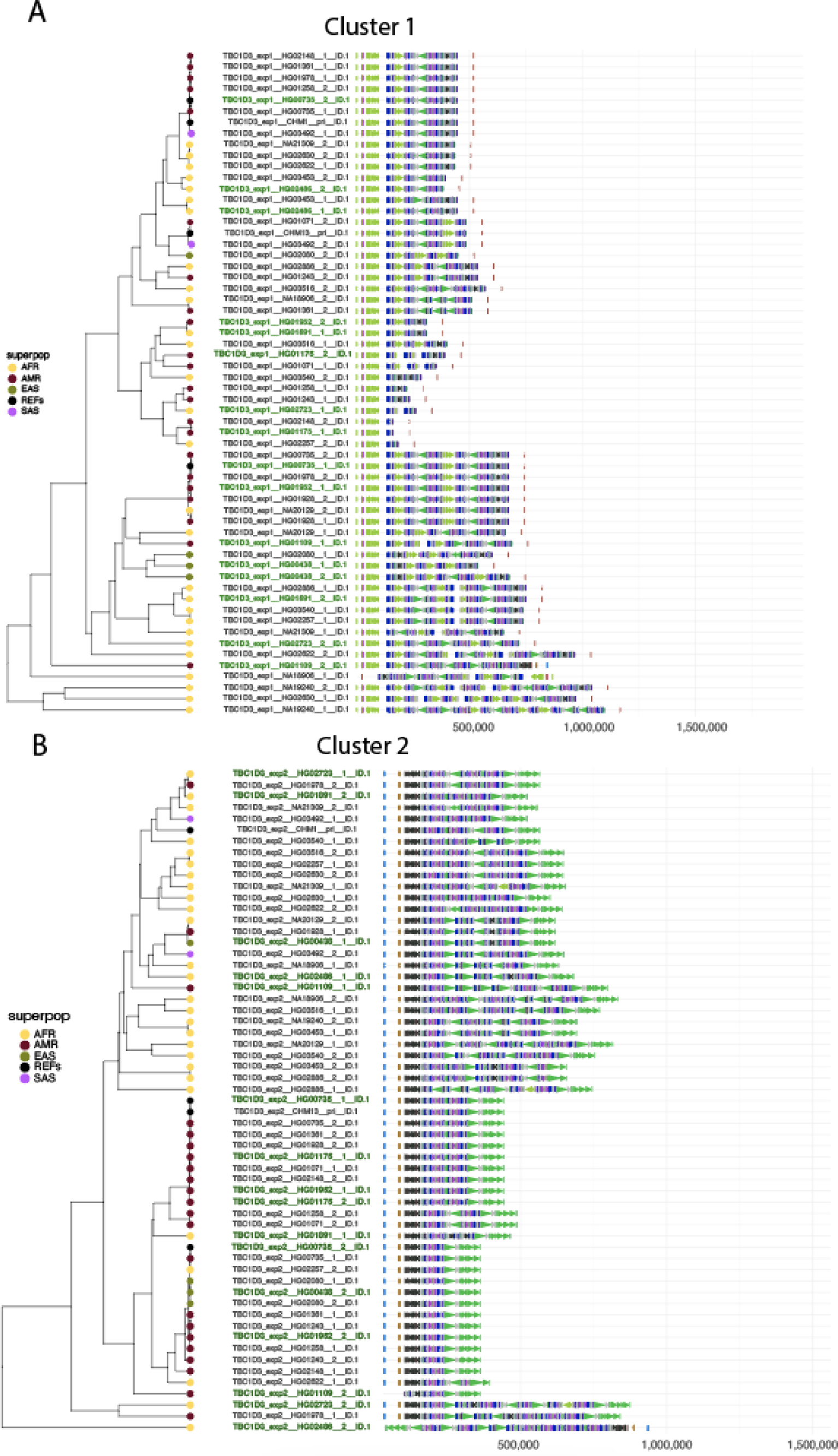
UPGMA clustering of *TBC1D3*. We hierarchically clustered validated assemblies based on duplicon content using UPGMA for cluster 1 **(A)** and 2 **(B)** (Methods). Superpopulations for each haplotype are included in the dendrogram, and assemblies rescued with Verkko are colored in green.

**Supplementary Figure S3:**
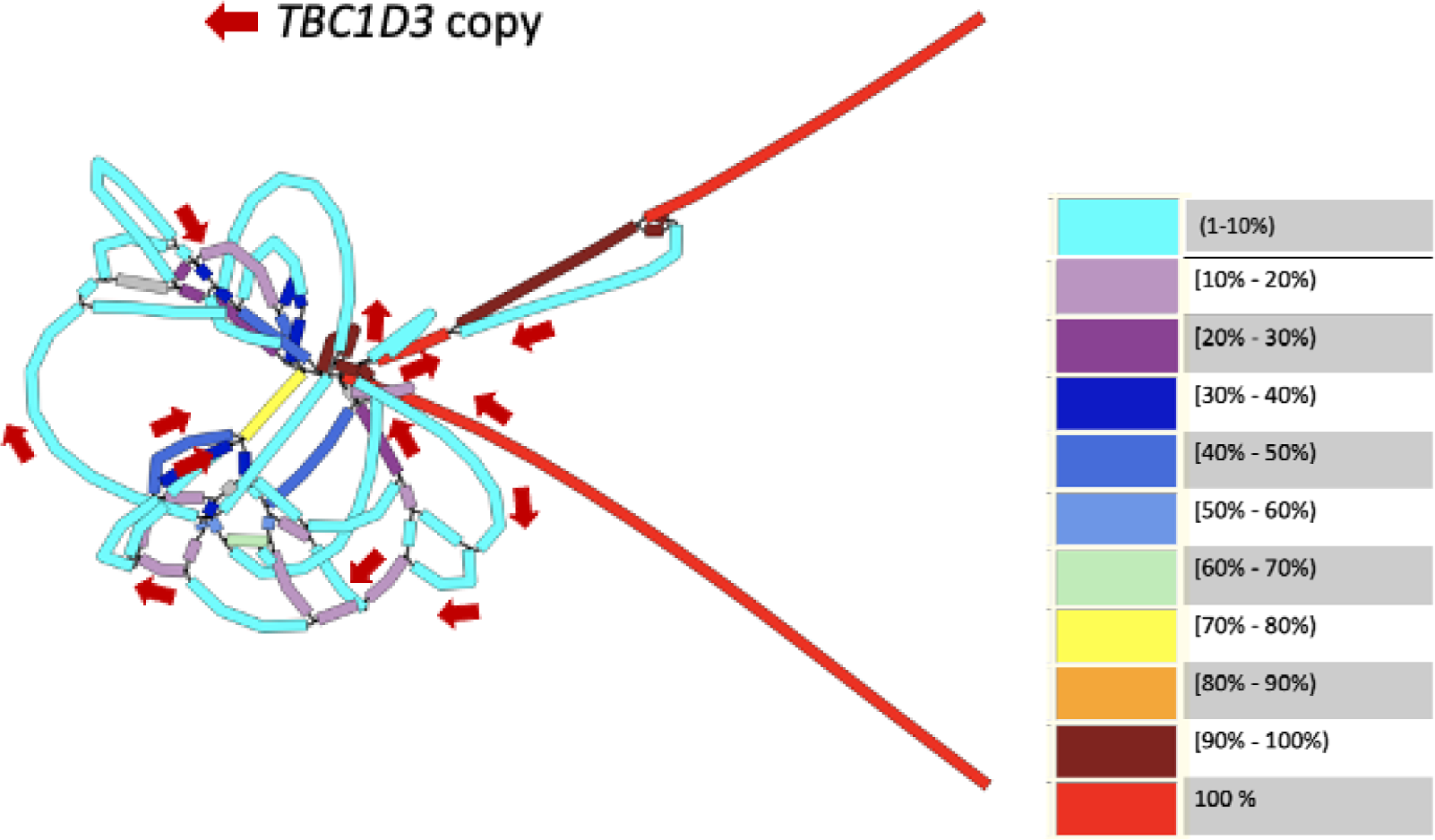
Minigraph pangenome graph. We generated a pangenome graph for *TBC1D3* using validated human haplotypes with Minigraph (settings -S -xggs -L 250 -r 100000). Graph segments are colored to represent the proportion of haplotypes that span the given segment, with light red indicating 100% representation and light blue indicating a single-haplotype traversal. *TBC1D3* paralogs are marked with arrows along the graph. We observe that *TBC1D3* structural variation is poorly reduced by Minigraph, where most copies reduce to nodes with single-haplotype support.

**Supplementary Figure S4:**
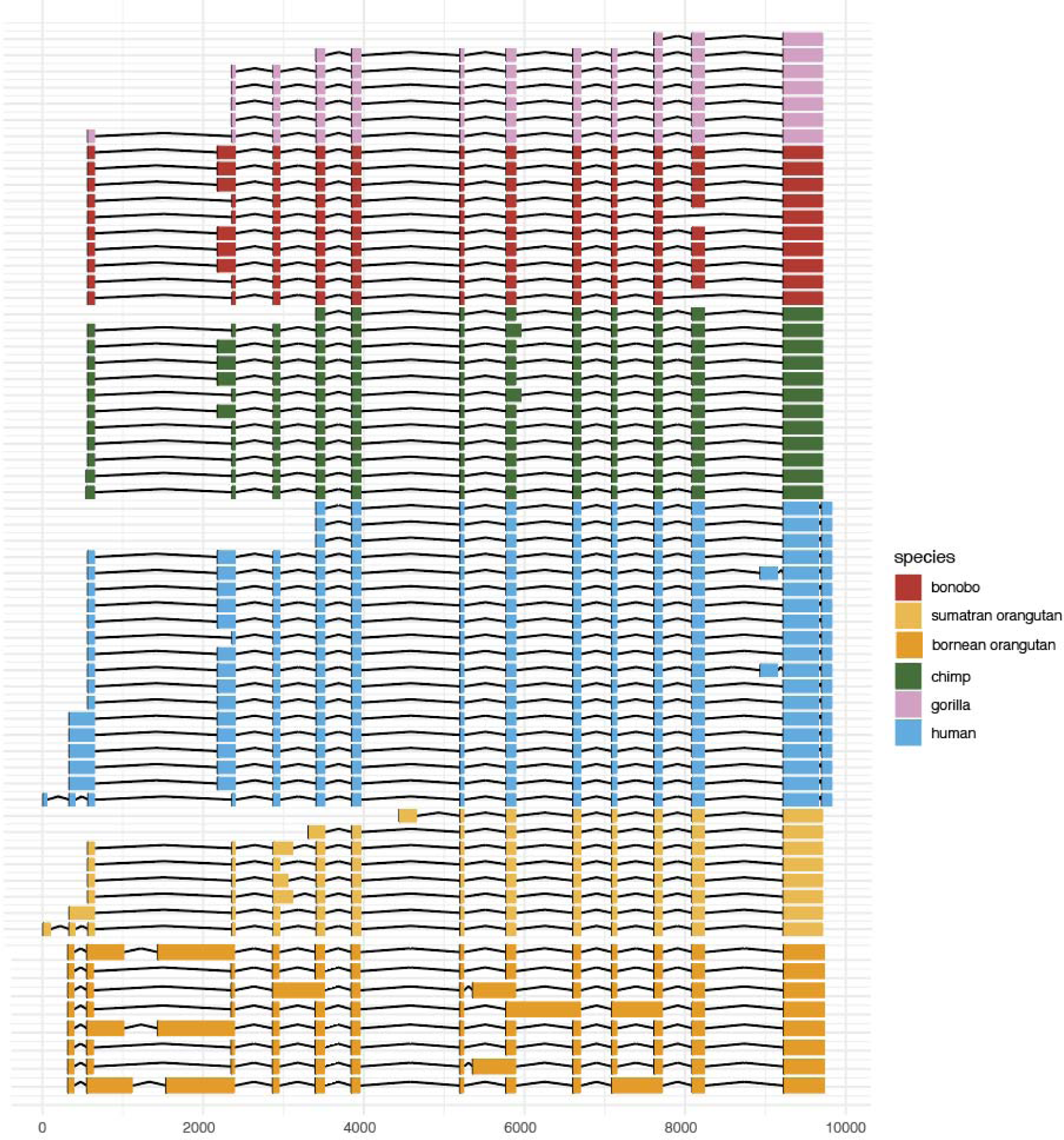
Expressed *TBC1D3* paralog isoforms. Paralog-specific isoforms were selected for each primate based on their length, mapping quality, and expression support. We observe expression of *TBC1D3* from all ape lineages examined; however, for ORF analysis Bornean orangutan and gorilla isoforms were removed due to lack of expression support or intron retention.

**Supplementary Figure S5:**
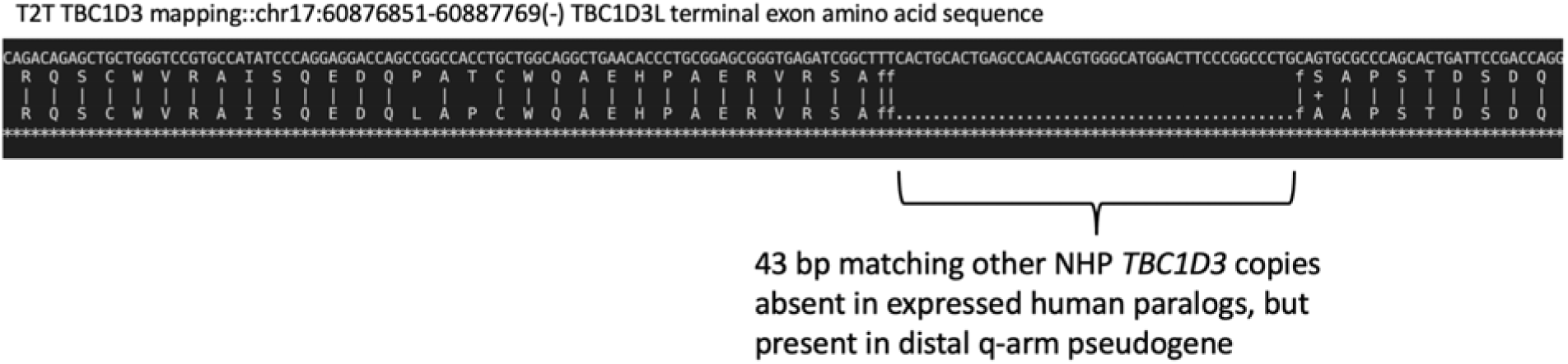
Missing 43 bp deletion in human distal *TBC1D3* pseudogenes. Human-expressed *TBC1D3*, with ORF modified by a 43 bp deletion, was mapped to all primate *TBC1D3* copies, including all human *TBC1D3* copies, with prosplign (Kiriyutin et al. 2017). Prosplign predicts the genomic nucleotides that represent codons of a given amino acid. We observe that distal *TBC1D3* pseudogenes lack the 43 bp deletion universally present in cluster 1 and 2 human *TBC1D3* copies.

**Supplementary Figure S6:**
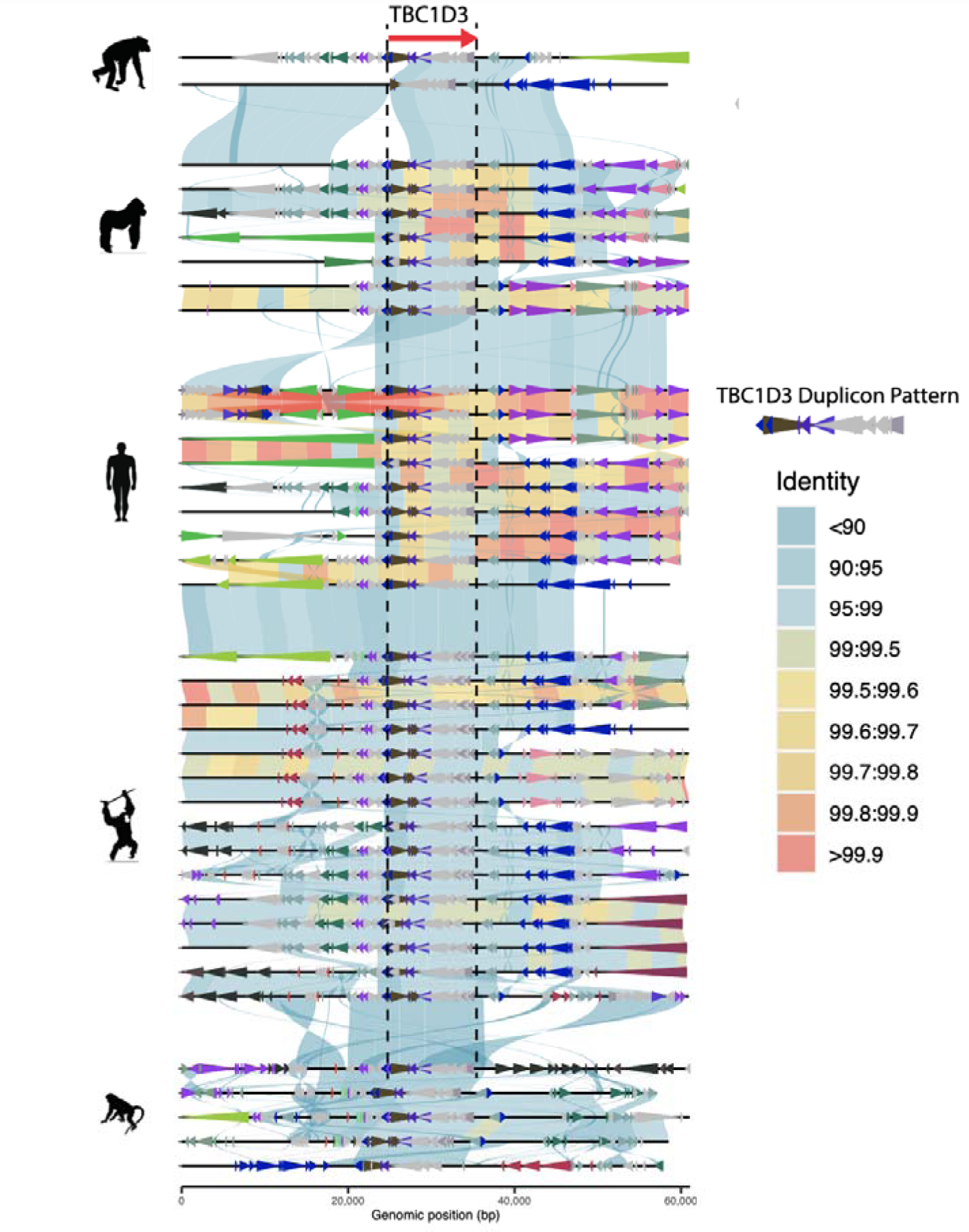
Local *TBC1D3* duplicon structure supports independent expansion. *TBC1D3* copies from clusters 1 and 2, along with 25 kbp flanking sequence, were extracted and mapped to one another. We observe that these paralogs consistently map best, with highest sequence identity and contiguity, to paralogs from their same species of origin, consistent with the hypothesis of independent expansion. This contiguity is similarly observed in underlying duplicon content, shown with colored arrows and annotated with DupMasker.

**Supplementary Figure S7:**
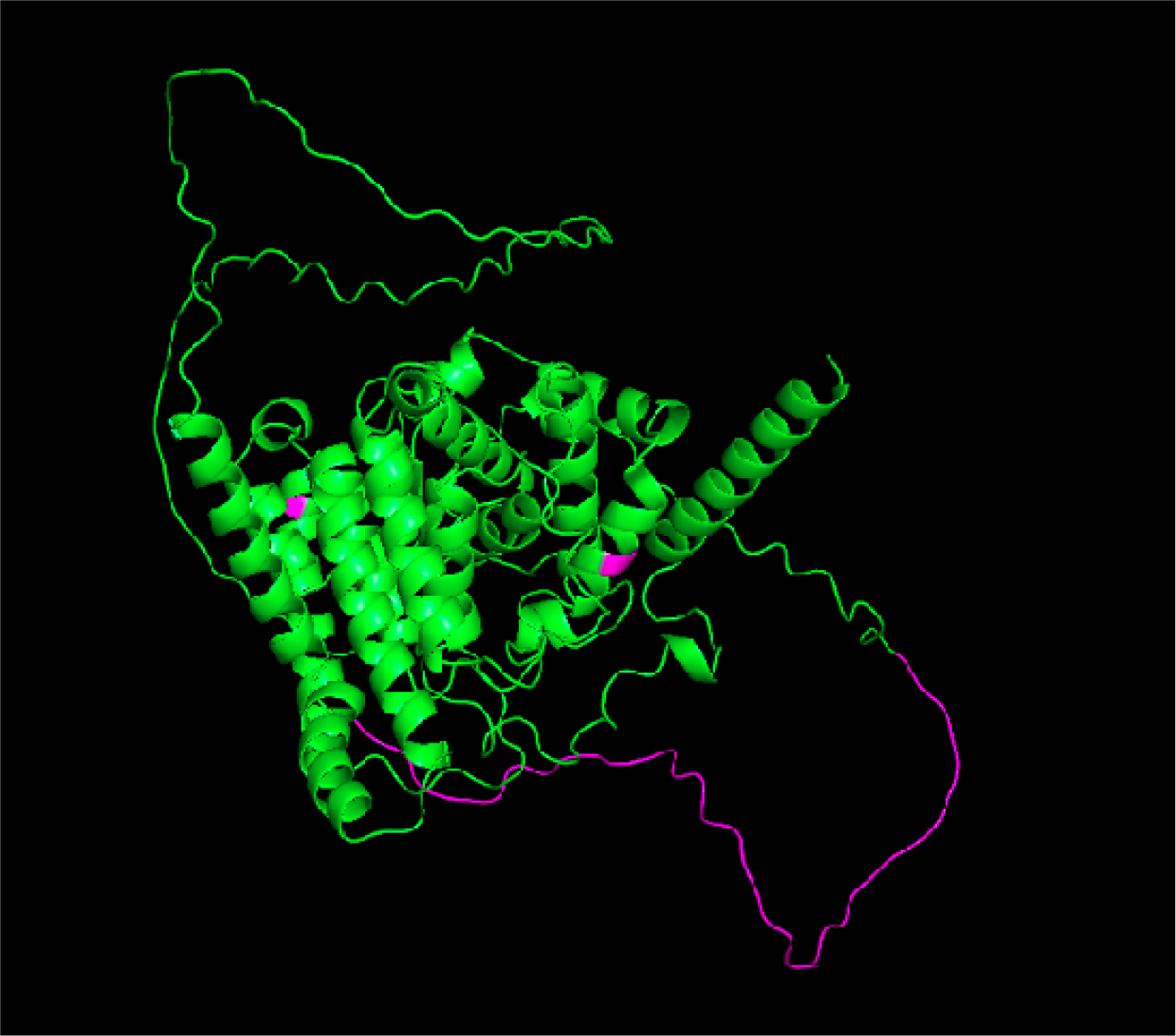
Human *TBC1D3* predicted tertiary structure. Human *TBC1D3* was predicted with Alphafold2 (https://alphafold.ebi.ac.uk/). Human lineage amino acid changes, including the modified carboxy terminus, are indicated with violet. We observe that the 41 aa novel C-terminus tertiary structure could not be predicted and is disordered.

**Supplementary Figure S8:**
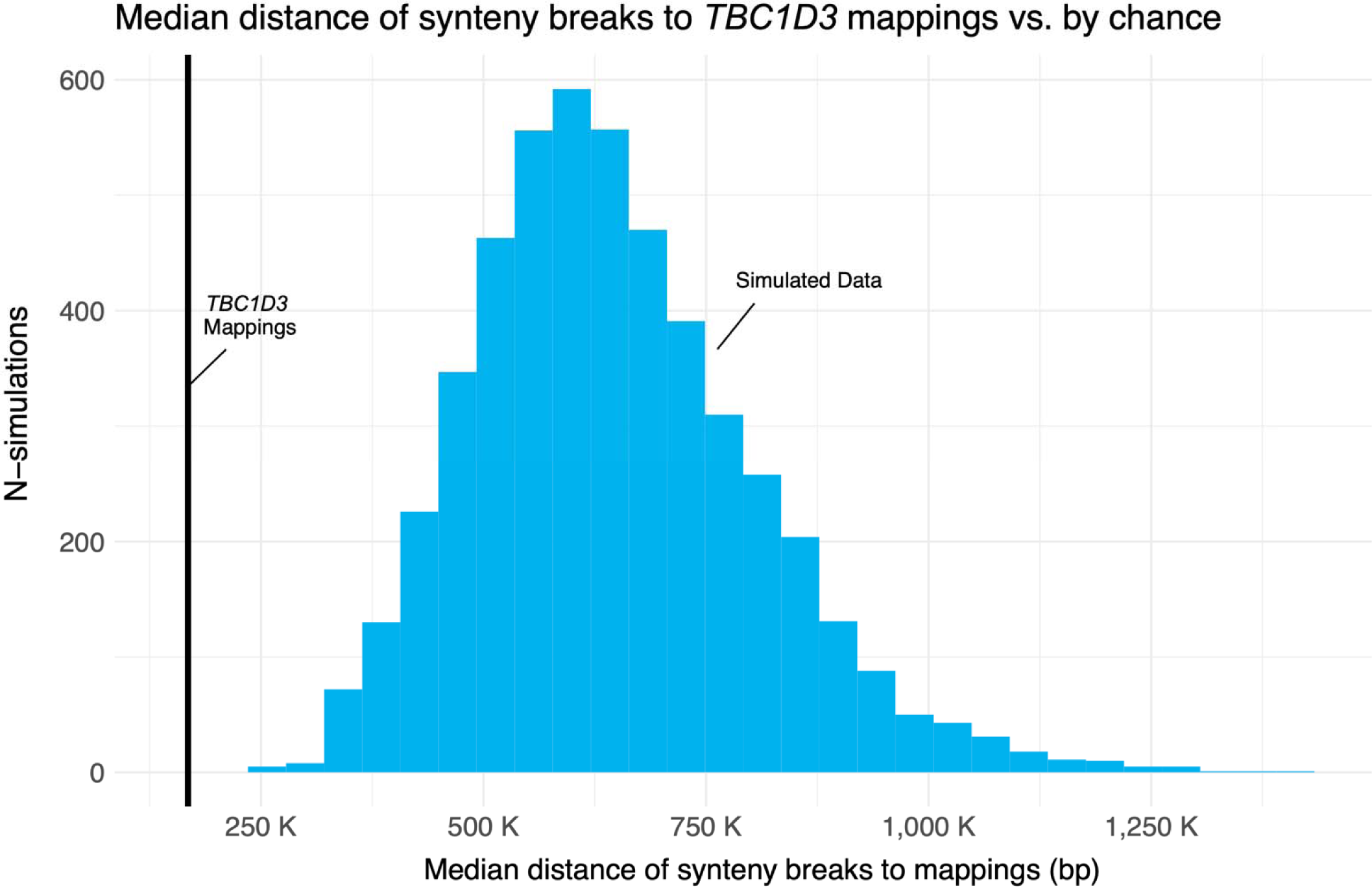
*TBC1D3* vs. random genomic sequence permutation. Sequences of 11 kbp were randomly selected from orthologous primate chromosome 17 contigs at the same quantity as the observed *TBC1D3* copies contained within the chromosome. We calculated the median distance of this sampling and repeated this experiment in 5000 permutations, comparing median distance relative to true *TBC1D3* mappings, marked in black.

**Supplementary Figure S9:**
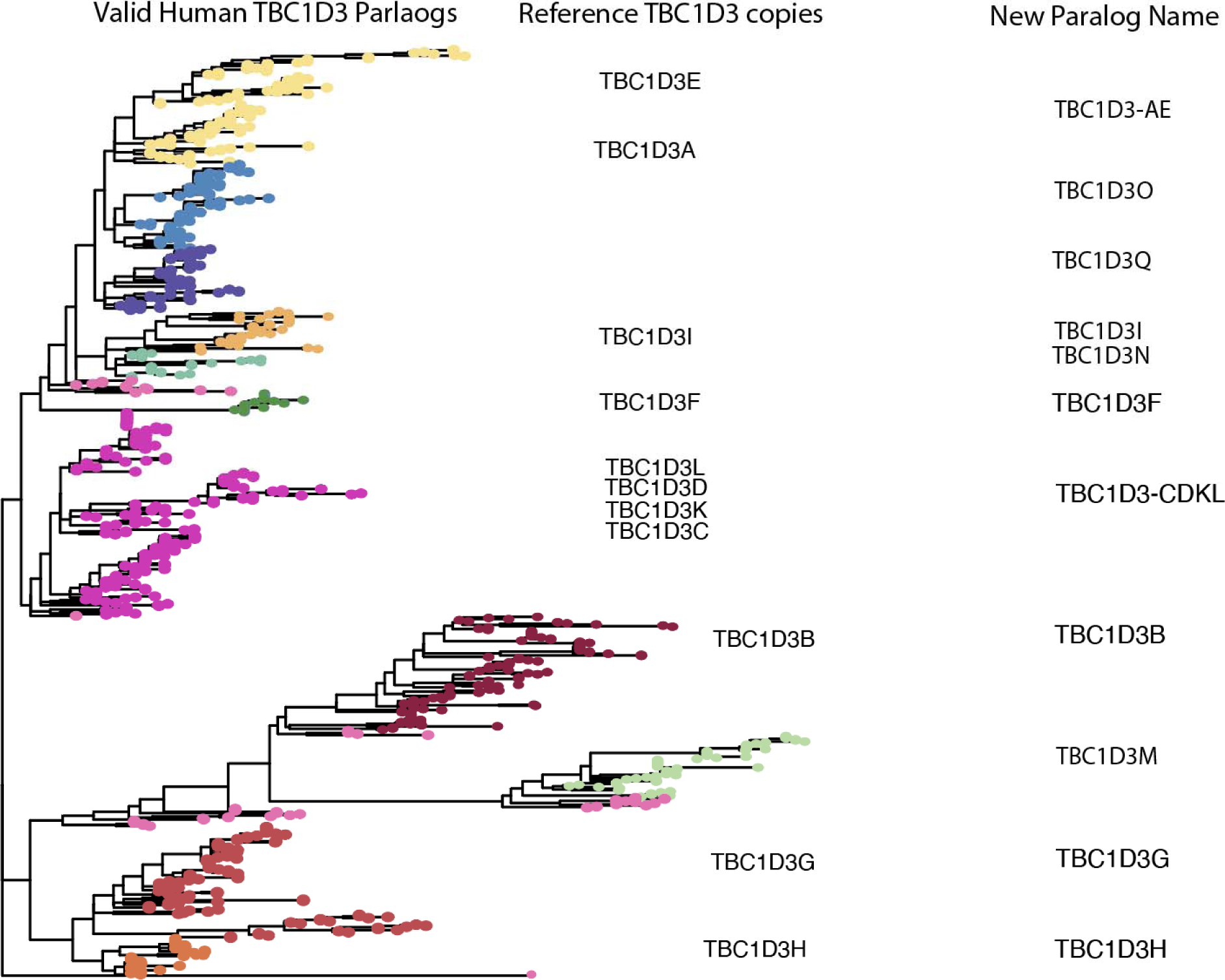
Pangenomic clustering and naming. Maximum likelihood phylogeny with ∼9600 bp of all human *TBC1D3* cluster 1 and 2 copies, outgrouped to chimpanzee *TBC1D3*. The paralog groups are defined by a heuristic intra-group allelic cutoff based on expected allelic variation in SD sequence (Methods). The first column of labels shows reference GRCh28 paralog locations within the phylogeny. The final column shows the new name given to the associated clusters. Most inherited the name assigned in GRCh28, or a concatenation when multiple paralogs mapped to a common cluster (*TBC1D3-AE; TBC1D3-CDKL*). Four novel population paralogs not included in GRCh38 were identified (*TBC1D3-M,TBC1D3-N, TBC1D3-O, TBC1D3-Q*).

**Supplementary Figure S10:**
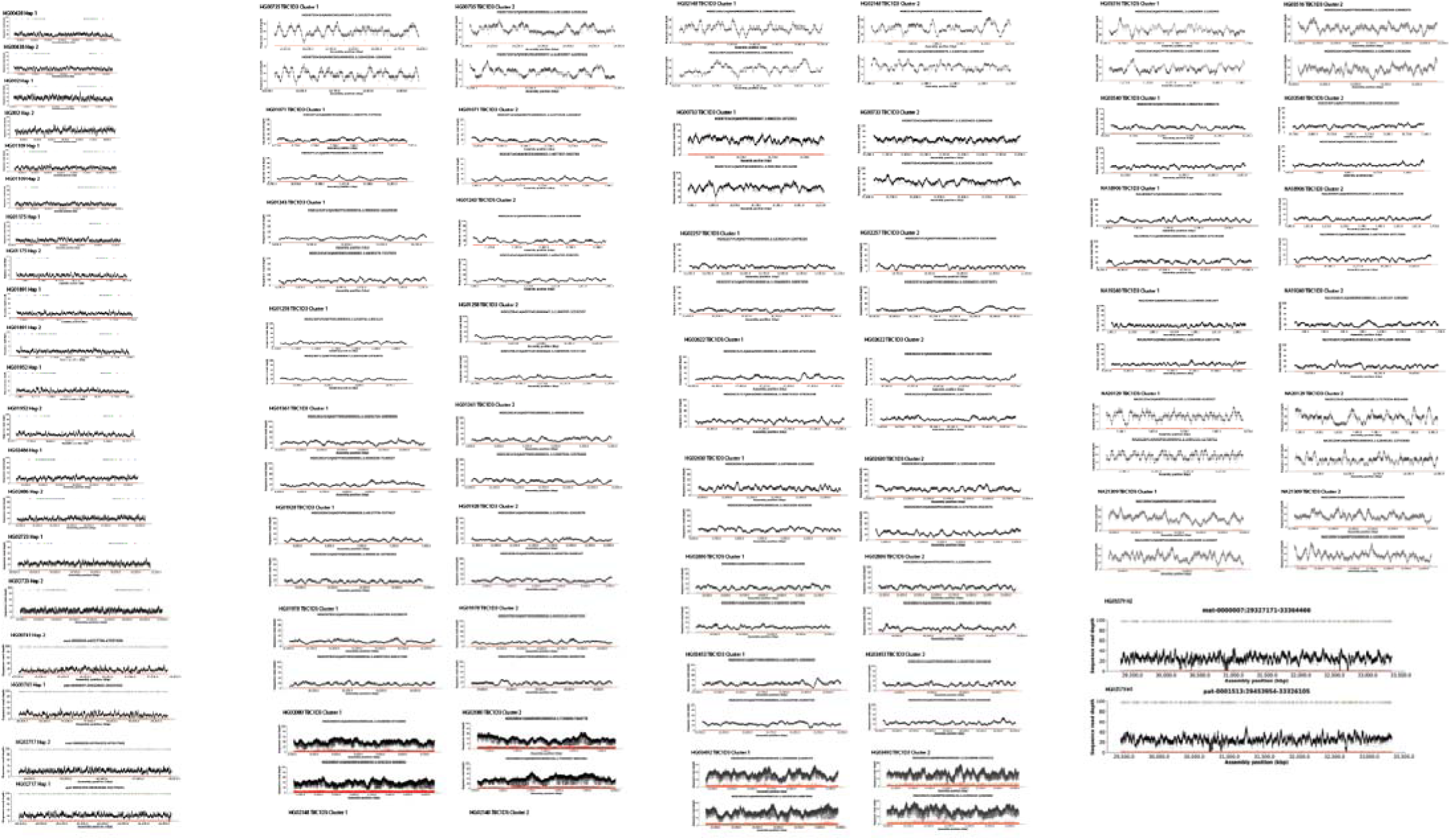
NucFreq validation. The haplotypes’ assembly accuracy was tested with NucFreq. Here, accuracy over each haplotype of *TBC1D3* cluster 1 and cluster 2 is demonstrated. Endogenous HiFi reads used to assemble respective phylogenies were mapped back onto the respective assemblies. We illustrated primary and secondary most prevalent base calls for reads aligned over a portion of an assembly in black and red, respectively. We expect primary base-call coverage to be relatively consistent, and consistently higher than the secondary base call, which should illustrate the sparse inaccuracies of HiFi sequencing.

## SUPPLEMENTARY TABLES

**Supplementary Table S1:** Primate *TBC1D3* copy number.

| Species | Scientific Name | Number of *TBC1D3* loci per haplotype \| | Number of *TBC1D3* genes (Hap1 \| Hap2) |
| --- | --- | --- | --- |
| Human *** | *Homo sapiens* | 4 \| 4 | 11 \| 22 |
| Gorilla* | *Gorilla gorilla* | 4 \| 4 | 13 \| 11 |
| Chimpanzee* | *Pan troglodytes* | 4 \| 4 | 8 \| 8 |
| Bonobo* | *Pan paniscus* | 4 \| 4 | 9 \| 9 |
| Sumatran orangutan* | *Pongo abelii* | 8 \| 9 | 23 \| 23 |
| Bornean orangutan* | *Pongo pygmaeus* | 5 \| 7 | 17 \| 21 |
| Gibbon* | *Symphalangus syndactylus* | 11 \| 11 | 31 \| 30 |
| Macaque** | *Macaca mulatta* | 4 \| 4 | 29 \| 20 |
| Gelada | *Theropithecus gelada* | 3 \| 3 | 31 \| 24 |
| Owl monkey** | *Aotus nancymaae* | 4 \| 4 | 8 \| 8 |
| Marmoset** | *Callithrix jacchus* | 2 \| 2 | 2 \| 2 |
| Mouse Lemur | *Microcebus murinus* | 0 \| 0 | 0 \| 0 |

\*Genomes sequenced as part of the Primate T2T Consortium (Makova et al. 2023). \*\*Genomes sequenced and assembled as part of Mao et al. 2024. \*\*\*Genomes sequenced as part of human T2T project and Human Pangenome Reference Consortium (Nurk et al. 2022; Liao et al. 2023).

**Supplementary Table S2:** Primate genome assembly statistics.

| Species | Scientific Name | HiFi Coverage | ONT Coverage | Assembly Method | Assembly N50 (Hap1 \| Hap2) (Mbp) |
| --- | --- | --- | --- | --- | --- |
| Gelada | *Theropithecus gelada* | 50.24 | NA | Hifiasm 0.15.2 | 109.0 \| 93.6 |
| Mouse Lemur | *Microcebus murinus* | 30.25 | NA | Hifiasm 0.15.2 | 29.4 \| 26.7 |
| Human | *Homo sapiens* | 30 | 120 | NA | 150.62 |
| Gorilla* | *Gorilla gorilla* | 107 | 165 | Verkko 1.4 | 151.43 \| 150.80 |
| Chimpanzee* | *Pan troglodytes* | 69 | 127 | Verkko 1.4 | 147.43 \| 140.84 |
| Bonobo* | *Pan paniscus* | 131 | 169 | Verkko 1.4 | 147.03 \| 147.48 |
| Sumatran orangutan* | *Pongo abelii* | 102 | 91.9 | Verkko 1.4 | 143.46 \| 140.60 |
| Bornean orangutan* | *Pongo pygmaeus* | 62.6 | 200 | Verkko 1.4 | 139.77 \| 139.21 |
| Gibbon* | *Symphalangus syndactylus* | 83.2 | 111 | Verkko 1.4 | 144.67 \| 145.50 |
| Macaque** | *Macaca mulatta* | 38.91 | NA | Hifiasm 0.15.2 | 18.81 \| 19.01 |
| Owl Monkey** | *Aotus nancymaae* | 36.57 | NA | Hifiasm 0.15.2 | 55.92 \| 44.99 |
| Marmoset** | *Callithrix jacchus* | 39.08 | NA | Hifiasm 0.15.2 | 103.97 \| 87.06 |

\*Genomes sequenced as part of the Primate T2T Consortium (Makova et al. 2023). \*\*Genomes sequenced and assembled as part of Mao et al. 2024.

**Supplementary Table S3:** Summary of Iso-Seq FLNC sequencing data.

| Sample | *TBC1D3* reference copies | Total number of sequencing reads | Reads mapped to *TBC1D3* |
| --- | --- | --- | --- |
| Human fetal brain | 14 | 2.33E+05 | 33138 |
| Human iPSCs** | 14 | 2.64E+08 | 11366 |
| Gorilla testis* | 15 | 4.68E+06 | 557 |
| Chimpanzee testis* | 8 | 3.00E+06 | 1461 |
| Bonobo testis* | 9 | 2.23E+06 | 915 |
| Sumatran orangutan testis* | 28 | 3.11E+06 | 2032 |
| Bornean orangutan testis* | 20 | 4.20E+06 | 6836 |

\*Iso-Seq FLNC data generated as part of Makova et al. 2023. \*\*Human iPSCs generated as part of Cheung et al. 2023.

**Supplementary Tables S4-S10**

Provided in a separate Excel file.

